# Calcium-dependent synaptic proteomics reveals EGFR signaling at active synapses

**DOI:** 10.64898/2026.05.20.726457

**Authors:** Adeline J. H. Yong, Yayu Wang, Juan A. Oses-Prieto, Tong Cheng, Chao Chen, Lane Domke, Mario V. Zubia, Angelina Koroknay, Jeffrey Simms, Alma L. Burlingame, Lily Y. Jan, Nicholas T. Ingolia, J. Wren Kim, Yuh Nung Jan

## Abstract

Activity-dependent remodeling of the synaptic proteome is essential for circuit maturation, plasticity, and adaptation. However, these molecular changes are transient and spatially sparse in intact neural circuits, making them difficult to resolve under physiological conditions. To address this challenge, we developed synaptic Cal-ID, a synapse-targeted, calcium-dependent proximity labeling strategy that enables activity-resolved proteomic profiling under native physiological conditions. Using synaptic Cal-ID, we uncover an unexpected mechanism by which synaptic activity locally engages epidermal growth factor receptor (EGFR) signaling. We identify two novel synaptic proteins, Anks1a and Ubash3b, that are rapidly recruited following activity and cooperatively promote postsynaptic enrichment and signaling of EGFR. Disruption of this pathway impairs synaptic maturation, plasticity, and memory. Together, these findings reveal how neuronal activity establishes local signaling competence at active synapses by concentrating the molecular machinery required to engage a broadly acting growth factor pathway.

## Introduction

Synaptic function relies on activity-dependent remodeling of protein composition to regulate synaptic strength, driving synaptic maturation and plasticity that shape neural circuit function^1–5^. Recent advances in proximity labeling-based proteomics have expanded the coverage of synaptic proteome profiling, revealing new molecular regulators of synapse assembly, function and plasticity^6–9^. However, these approaches primarily capture steady-state composition and cannot selectively resolve proteins associated with synaptic activity, limiting our ability to define how synaptic proteomes are remodeled by activity. This challenge arises because activity occurs transiently across a sparse subset of synapses at any given time, such that proteins enriched during these events comprise only a small fraction of the total synaptic proteome and are further masked by the preferential detection of high-abundance proteins by mass spectrometry^10^. To circumvent this, most studies have relied on experimental stimulation to broadly activate synapses, rather than capturing changes arising from natural patterns of activity.

Calcium is a central signaling molecule in excitatory synapses. Activation of postsynaptic glutamate receptors, particularly N-methyl-D-aspartate receptors (NMDARs), generates transient calcium elevations that, above a threshold, engage calcium-binding proteins to regulate synaptic structure and function^11–14^. While postsynaptically targeted calcium indicators, including PSD-95-tagged GCaMP^15^ and PSD-95-tagged CAMPARI (SynTagMA)^16^, have provided valuable insights into the timing and spatial distribution of calcium signals at individual synapses, they cannot reveal the underlying molecular events associated with these signals.

We recently developed Cal-ID, a calcium-dependent promiscuous biotin ligase that becomes enzymatically active upon calcium-induced conformational refolding^17^, enabling labelling and identification of proteins associated with calcium-dependent processes. Importantly, Cal-ID responds to physiological calcium dynamics within cultured neurons and brains^17^. Given that synaptic activity drives local calcium elevations, we developed a synapse-targeted Cal-ID to selectively label proteins within active synapses. This approach is particularly valuable in the brain, where activity is restricted to specific circuits and subsets of synapses. We hypothesized that profiling active synapses would enable the capture of proteins that are selectively enriched and/or transiently recruited, thereby uncovering new regulators of activity-dependent synaptic remodeling.

### A calcium-dependent biotin ligase for activity-dependent labeling at synapses

We engineered synaptic Cal-ID for activity-dependent postsynaptic labeling at synapses (Fig. 1c) by tagging Cal-ID with PSD-95 (Syn-Cal-ID; Fig. 1a–b), a major excitatory postsynaptic scaffolding protein that has previously been used to target the proximity labelling enzyme BioID to synapses^7^. We generated cpTurboID-PSD95 (Syn-cpTurboID; Fig. 1a–b) as a control for Syn-Cal-ID by removing its calcium-responsive domains. When we expressed Syn-Cal-ID and Syn-cpTurboID in neurons, V5 and biotinylation signals overlap with the AMPAR subunit GluA1, consistent with enzyme enrichment and biotinylation of proteins at the postsynaptic density (PSD) (Fig. 1d and Extended Data Fig. 1a–c). To validate that Syn-Cal-ID responds to synaptic calcium increase, we induced chemical long-term potentiation (cLTP), a protocol that activates glutamate receptors and elevates synaptic calcium^18–20^, in primary mouse cortical neurons. Neurons expressing Syn-Cal-ID showed increased protein biotinylation at synaptic regions following cLTP (Fig. 1e–f and Extended Data Fig. 1d–e), whereas this increase was absent in the Syn-cpTurboID control (Fig. 1g and Extended Data Fig. 1f). To test Syn-Cal-ID’s response to reduced synaptic calcium, we treated neurons with either BAPTA-AM, a membrane-permeable calcium chelator, or a cocktail of drugs to suppress postsynaptic calcium entry by inhibiting ionotropic glutamate receptors and voltage-gated calcium channels (Extended Data Fig. 1g). We observed decreased biotinylation of PSD proteins in Syn-Cal-ID expressing neurons treated with BAPTA-AM or the drug cocktail, compared to untreated neurons (Fig. 1h). Meanwhile, PSD protein biotinylation in Syn-cpTurboID expressing neurons remained similar in all conditions (Fig. 1h). Collectively, these experiments indicate that Syn-Cal-ID can respond dynamically to changes in synaptic calcium levels.

**Figure 1.**
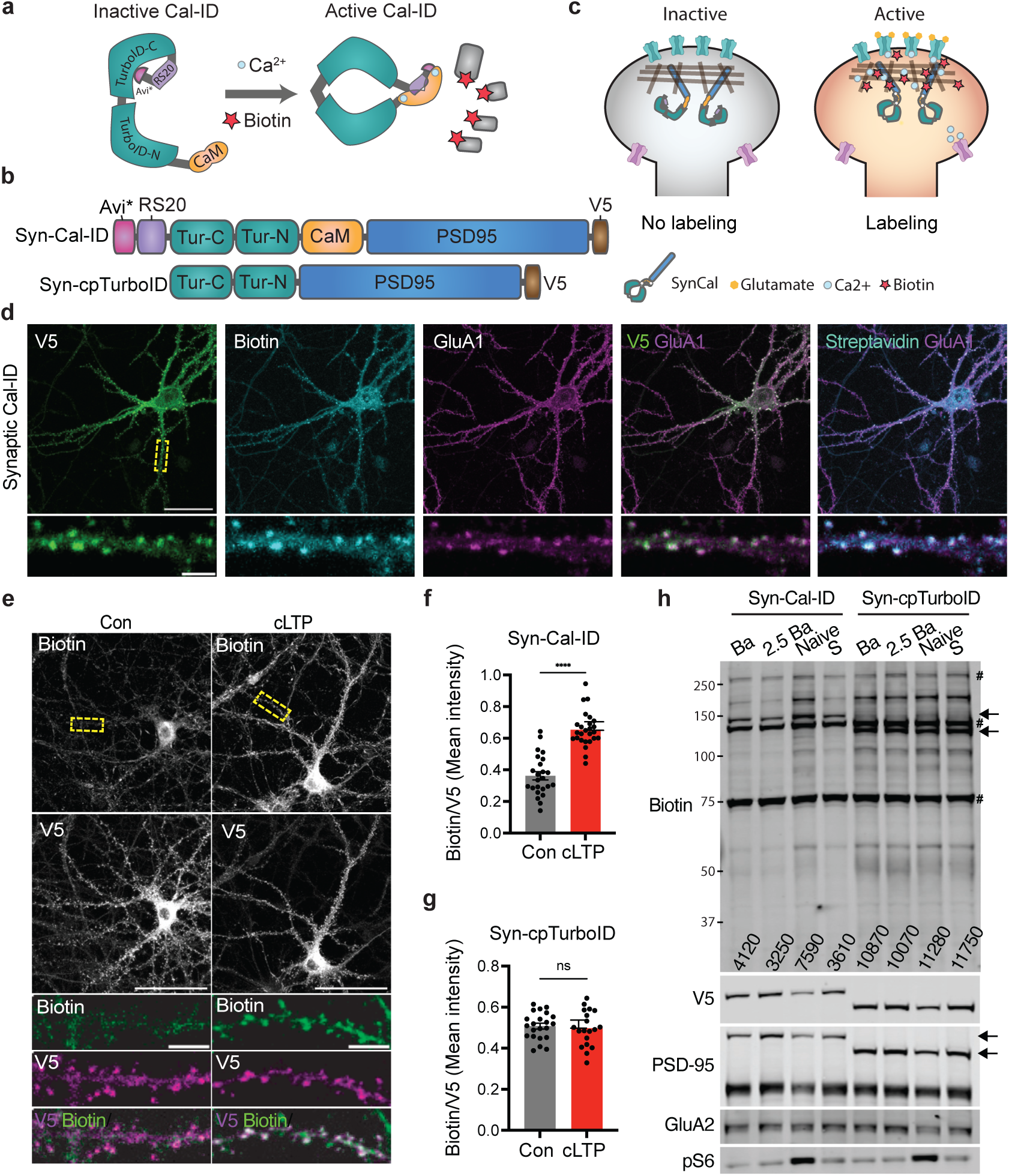
Synaptic Cal-ID is a synapse-localized calcium-dependent biotin ligase. a, Schematic of Cal-ID activation. Calcium-dependent activation of circularly permutated TurboID. b, Protein domains for Syn-Cal-ID and Syn-cpTurboID. Tur-C, TurboID C-terminus. Tur-N, TurboID N-terminus. c, Syn-Cal-ID labelling of proteins within active synapses. d, Immunostaining of primary mouse neurons transduced with lentivirus expressing Syn-Cal-ID. e–g, representative images (e) and quantification (f, g) of mouse cultured neurons transfected with Syn-Cal-ID (e, f) or Syn-cpTurboID (g) following chemical LTP (cLTP) or control. F, Welch’s t-test, *n* = 24–26 cells, 4 culture batches. g, Welch’s t-test, *n* = 19–22 cells, 4 culture batches. h, Western blot analysis of PSD fractions from primary mouse neurons transduced with lentivirus expressing Syn-Cal-ID or Syn-cpTurboID and treated with 40uM (Ba), or 100uM BAPTA-Am (2.5Ba), or silenced with 50μM APV, 20μM NBQX, 20μM Cilnepdine, 10μM Nifedipine, 20μM Mifepadril (S), or left untreated. #: endogenous biotinylated proteins. Arrows: Syn-Cal-ID or Syn-cpTurboID. DIV19–21 primary mouse cortical neuron cultures. 100uM biotin labelling for 20 min (e–g) or 1 h (d, h). Scale bars 50μm or 5μm (zoom). Data are presented as mean ± s.e.m. ns, not significant. ****P < 0.0001.

### Activity-dependent synaptic proteomics from cultured neurons reveal Ubash3b as a novel activity-dependent synaptic protein

To obtain a more comprehensive overview of the activity-dependent synaptic proteome, we chose to perform proteomic analysis from both cultured mouse neurons and the mouse brain. Cultured neurons enable assessment of cell-autonomous synaptic responses, revealing how individual synapses intrinsically respond to activity, whereas neurons in the brain provide a richer circuit context, reflecting how neurons are connected and influenced by non-cell-autonomous factors such as glia. Together, these complementary approaches provide a broader and more physiologically relevant view of activity-dependent synaptic remodeling.

Given the high spontaneous activity of cultured neurons^21,22^, we did not apply external stimulation. Instead, Syn-Cal-ID labeling was performed during baseline activity to capture synapses associated with ongoing neuronal activity. We transduced cultured neurons with lentivirus containing Syn-Cal-ID or Syn-cpTurboID or left them untreated as a control. Syn-Cal-ID selectively labels proteins at active synapses, whereas Syn-cpTurboID labels proteins at all synapses, providing a measure of baseline (average) synaptic composition. Untransduced neurons served as a control for non-specific binding during streptavidin pulldown (Fig. 2a). Neurons were treated with biotin, followed by subcellular fractionation to isolate PSD proteins and streptavidin pulldown to enrich for biotinylated PSD proteins (Fig. 2a). Western blot analysis of the biotinylated PSD proteins showed that the PSD proteins, including PSD-95, AMPAR subunit GluA2, and Homer were biotinylated in both Syn-Cal-ID- and Syn-cpTurboID-expressing neurons, but not in the no virus control (Fig. 2b). Notably, Syn-Cal-ID labelling was readily detected under untreated conditions, indicating that baseline synaptic activity is sufficient to drive activity-dependent protein labeling. Furthermore, streptavidin bead pulldown was effective and captured most biotinylated proteins (Extended Data Fig. 2a). We next performed TMT (tandem mass tag)-based quantitative mass spectrometry and identified 1515 proteins represented by 9498 unique peptides. Principal component analysis (PCA) showed that samples cluster according to condition (Extended Data Fig. 2b), indicating good reproducibility and separation between groups. Following normalization (Methods), nonlinear regression revealed an overall positive relationship between baseline synaptic abundance (Syn-cpTurboID) and Syn-Cal-ID labeling. Proteins were then ranked based on the deviation between observed Syn-Cal-ID labeling and the labeling predicted from this regression trend, with positive deviations indicating greater-than-expected enrichment at active synapses (Supplementary Table 1). Calmodulin showed the highest enrichment ratio (Fig. 2c), which is likely due to self-biotinylation of Syn-Cal-ID^17^. As expected, PSD-95 shows no enrichment, as its presence in both Syn-Cal-ID and Syn-cpTurboID constructs leads to comparable self-labeling by the two enzymes (Fig. 2c). To characterize top enriched targets, proteins with a Z-score greater than 1.3 (top 4.7%) were selected for further analysis. These include GluA3, BRAG2^23^, CTNND2^24^ and UCHL1^25^, which have known functions in synaptic plasticity and/or learning and memory (Fig. 2c), supporting the ability of this strategy to identify activity-related synaptic proteins. Furthermore, STRING network analysis identified three clusters of proteins associated with modulation of chemical synaptic transmission, translation, and intracellular transport, indicating functional coupling of activity-enriched proteins to synaptic signaling, protein synthesis, and trafficking (Extended Data Fig. 2c).

**Figure 2.**
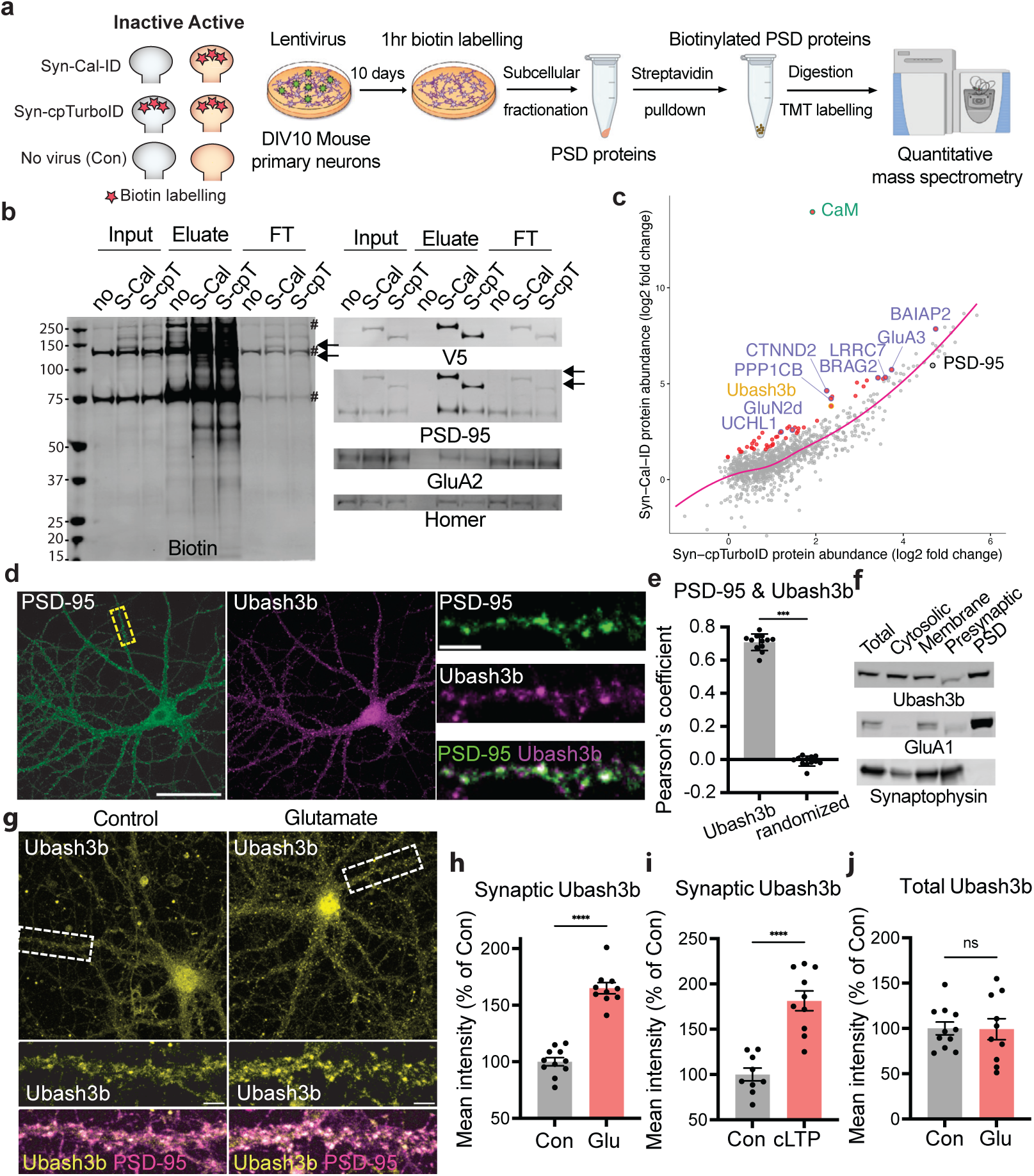
Proteomic screen using synaptic Cal-ID in vitro identifies Ubash3b as a novel activity-dependent synaptic protein. a, Experimental strategy. b, Western blot of PSD fraction after streptavidin pulldown (input, eluate or flow-through [FT]) from neurons expressing no virus (no), Syn-Cal-ID (S-Cal) or Syn-cpTurboID (S-cpT). #: endogenous biotinylated proteins. Arrows: Syn-Cal-ID or Syn-cpTurboID. c, Mass spectrometry-based relative protein abundance (normalized to no virus) comparing Syn-Cal-ID and Syn-cpTurboID conditions. Pink line: local polynomial regression using LOESS. Purple: known synaptic plasticity-related proteins. Red dots: proteins above 1.64 Z-score cut off. d–e, Representative image (d) and quantification (e) of immunostaining. e, Wilcoxin matched-pairs signed rank test, *n* = 12 cells, 2 batches of cultures. F, Western blot from subcellular fractionation of 5-month-old mouse cortex. g-j, Representative images (g) and quantifications (h–j) of immunostaining of mouse cultured neurons treated with 10uM glutamate for 5mins (g,h,j) or chemical LTP (cLTP; i). Synaptic Ubash3b is defined as Ubash3b signal within PSD-95 puncta. h, j, Mann-Whitney test, *n* = 10–11 cells, 2 culture batches. i, Mann-Whitney test, *n* = 9–10, 3 culture batches. DIV19–21 primary mouse cortical neuron cultures. Scale bars 50μm (whole cell) or 5μm (zoom). Data are presented as mean ± s.e.m. ns, not significant. ***P < 0.001, ****P < 0.0001.

Alongside known activity-associated proteins identified, we identified several proteins not previously characterized at synapses, possibly representing low-abundance, dynamically recruited regulators of synaptic remodeling that were missed by prior proteomic approaches. From these, we prioritized Ubash3b for further study given its known role as an atypical tyrosine phosphatase that regulates receptor-mediated signaling, including EGFR signaling^26,27^, suggesting the potential to influence multiple synaptic processes. We first examined the neuronal localization of Ubash3b. Immunostainings of cultured mouse neurons (Fig. 2d–e) and western blot from PSD fractions of the adult mouse cortex (Fig. 2f) both revealed synaptic localization of Ubash3b. Because our calcium-dependent proteomic screen suggests that Ubash3b is enriched at active synapses, we tested whether synaptic activity modulates its localization. Increasing synaptic activity with glutamate^28^ or cLTP increased Ubash3b in synaptic regions (Fig. 2g–i and Extended Data Fig. 2d), without changing total protein levels (Fig. 2j and Extended Data Fig. 2e), indicating that Ubash3b is a novel synaptic protein that undergoes activity-dependent synaptic enrichment.

### Activity-dependent synaptic proteomics from cortex reveal Anks1a as novel activity-dependent synaptic protein

To ensure sufficient Syn-Cal-ID labelling in the brain, we used enriched environment (EE; Extended Data Fig. 3e), a behavior widely used in rodents to study naturally occurring changes in the brain during a learning and memory-related experience. Among other effects, EE results in brain-wide increases in cell excitability, synapse number and synaptic strength^29,30^ without imposing stress to the animal. We injected adeno-associated viruses (AAVs) expressing Syn-Cal-ID or Syn-cpTurboID under the human Synapsin promoter bilaterally into the cortex of postnatal day 0 (P0) mice pups and included a no-virus control for comparison (Fig. 3a–b and Extended Fig. 3a). After 4–5 months, we administered biotin and immediately placed the mice into EE for 4 hours, a duration which allows for sufficient biotinylation in synapses (Fig. 3c and Extended data Fig. 3c). PSD fractions were isolated from mouse cortices and biotinylated PSD proteins captured using streptavidin beads (Extended Data Fig. 3b, 3d), followed by TMT-based quantitative mass spectrometry. 4865 peptides from 608 proteins were identified. PCA showed consistent clustering of samples (Extended Data Fig. 4a), supporting reproducibility. We calculated enrichment ratios of each protein similarly to the screen from cultured neurons (Fig. 3d) and ranked the proteins (Supplementary Table 1). As in our in vitro screen, the top hit was Calmodulin, whereas PSD-95 showed no enrichment (Fig. 3d). Furthermore, SynGAP, a protein known to disperse from the PSD during activity^31^, was identified in the de-enriched fraction (Fig. 3d) consistent with its reduced presence at active synapses. Using a Z-score of 1, we selected the top 7.6% proteins for further analysis. These included BRAG2^23^, SAP97^32^, SHISA6^33^ and SAPAP4^34^, all of which have established roles in synaptic plasticity. STRING network analysis revealed enrichment of protein networks involved in modulation of chemical synaptic transmission and cytoskeletal organization (Extended Data Fig. 4b), indicating that proteins enriched at active synapses are functionally linked to synaptic signaling and structural plasticity.

**Figure 3.**
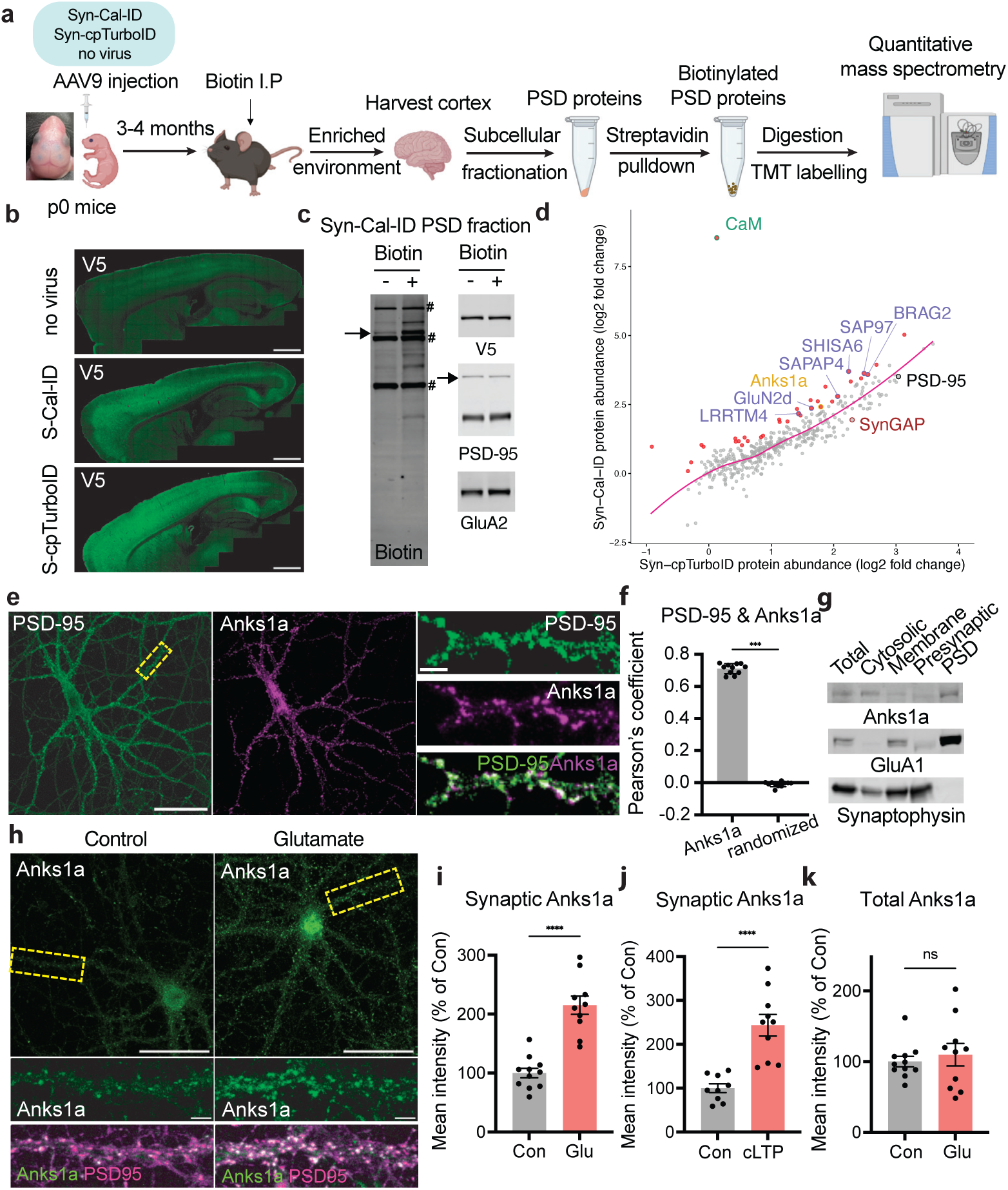
Proteomic screen using synaptic Cal-ID in vivo identifies Anks1a as a novel activity-dependent synaptic protein. a, Experimental strategy. b, Representative images of brains showing virus spread. c, Western blot of PSD fractions after streptavidin pulldown from mouse cortex expressing Syn-Cal-ID with or without biotin injection. #: endogenous biotinylated proteins. Arrows: Syn-Cal-ID. d, Representative images of brains of mice. e, Scatter plot of log₂ fold changes (relative to control) comparing Syn-Cal-ID and Syn-cpTurboID. The pink line shows a LOESS fit. Synaptic plasticity-related proteins are highlighted in purple, proteins above a Z-score cutoff of 1 in red. e–f, Representative image (e) and quantification (f) of immunostaining. Wilcoxin matched-pairs signed rank test, *n* = 11 cells, 3 batches of cultures. g, Western blot from subcellular fractionation of 5-month-old mouse cortex. H-K, Representative images (h) and quantifications (i–k) of immunostaining of mouse cultured neurons treated with 15uM glutamate for 5 mins (i, k) or chemical LTP (cLTP; j). Synaptic Anks1a is defined as Anks1a signal within PSD-95 puncta. I, K, Mann-Whitney test, *n* = 10–11 cells, 2 culture batches. j, Mann-Whitney test, *n* = 9–10, 3 culture batches. DIV19–21 primary mouse cortical neuron cultures. Scale bars 1mm (B), 50μm (whole cell) or 5μm (zoom). Data are presented as mean ± s.e.m. ns, not significant. ***P < 0.001, ****P < 0.0001.

While many top-ranked proteins had established synaptic roles, we next focused on Anks1a, a protein not previously characterized at synapses. Anks1a is an adaptor protein known to enhance receptor tyrosine kinase trafficking, including EGFR recycling to the cell surface while limiting its lysosomal degradation^35^. Notably, its function parallels that of Ubash3b, suggesting a shared role in promoting EGFR signaling at synapses. Immunostaining of cultured mouse neurons (Fig. 3e–f and Extended Data Fig. 4c) and western blot from PSD proteins in the adult mouse cortex (Fig. 3g) confirmed the localization of Anks1a to the PSD. Furthermore, increasing synaptic activity with glutamate or cLTP increased synaptic localization of Anks1a (Fig. 3h–j and Extended Data Fig. 4d) without altering total protein levels (Fig. 3k and Extended Data Fig. 4e), indicating that Anks1a is a synaptic protein recruited to synapses in response to activity.

### Mechanism of activity-dependent enrichment of Anks1a and Ubash3b at synapses

To define how rapidly Ank1a and Ubash3b become enriched at synapses, we performed a time course experiment and found that Anks1a and Ubash3b localize to synaptic regions within 1 minute of glutamate stimulation (Extended Data Fig. 5). Furthermore, their enrichment persisted in the presence of the translation inhibitor anisomycin (Extended Data Fig. 6a–c), indicating that enrichment is independent of new protein synthesis. However, it was abolished by NMDAR blockade with APV (Extended Data Fig. 6d–f), demonstrating NMDAR dependence. Moreover, glutamate stimulation in the presence of tetrodotoxin, which blocks action potentials, was sufficient to drive synaptic recruitment (Extended Data Fig. 6g–i), indicating that glutamate receptor activation alone is sufficient. Collectively, these findings reveal that Anks1a and Ubash3b are rapidly recruited to the PSD upon synaptic activation, and that this recruitment depends on NMDAR signaling but not protein synthesis.

### Anks1a and Ubash3b are required for synaptic maturation

Given the established role of neuronal activity in driving synaptic maturation^36–38^, we hypothesized that the activity-dependent recruitment of Anks1a and Ubash3b promotes synaptic maturation. To test this, we performed CRISPRi-mediated knockdown (KD) of each protein (∼95% efficiency; Extended data Fig. 7a–f) and assessed dendritic spine morphology, a proxy for synaptic strength^39^. Sparse transfection with a knockdown construct co-expressing mCherry enabled visualization of dendritic spines by cell fill. We observed differences in spine morphology in Anks1a-KD and Ubash3b-KD neurons, characterized by a reduced proportion of mushroom spines and increased proportions of long and filopodia-like spines compared to control neurons (Fig. 4a, c), without affecting spine density (Extended Data Fig. 7g). These changes are consistent with a shift toward a less mature spine phenotype. Furthermore, we stained for surface AMPARs, which are correlates of synaptic strength^3,12^ and observed decreases in both surface GluA1 (Fig. 4b, d) and GluA2 (Fig. 4e and Extended Data Fig. 7h) in Anks1a-KD and Ubash3b-KD neurons, indicating reduced synaptic strength. Consistently, PSD-95 staining revealed smaller and less intense puncta (Fig. 4f–h), indicating reduced synapse size, without a change in puncta density (Extended Data Fig. 7i), indicating no effect on synapse numbers. Finally, we directly assessed synapse function using miniature excitatory postsynaptic current (mEPSC) recordings. mEPSC analysis of Anks1a-KD and Ubash3b-KD neurons showed a decreased amplitude, with no change in frequency (Fig. 4i–l), consistent with weaker synapses without altered synapse number. Taken together, these findings indicate that Anks1a and Ubash3b are required for structural and functional maturation of synapses.

**Figure 4.**
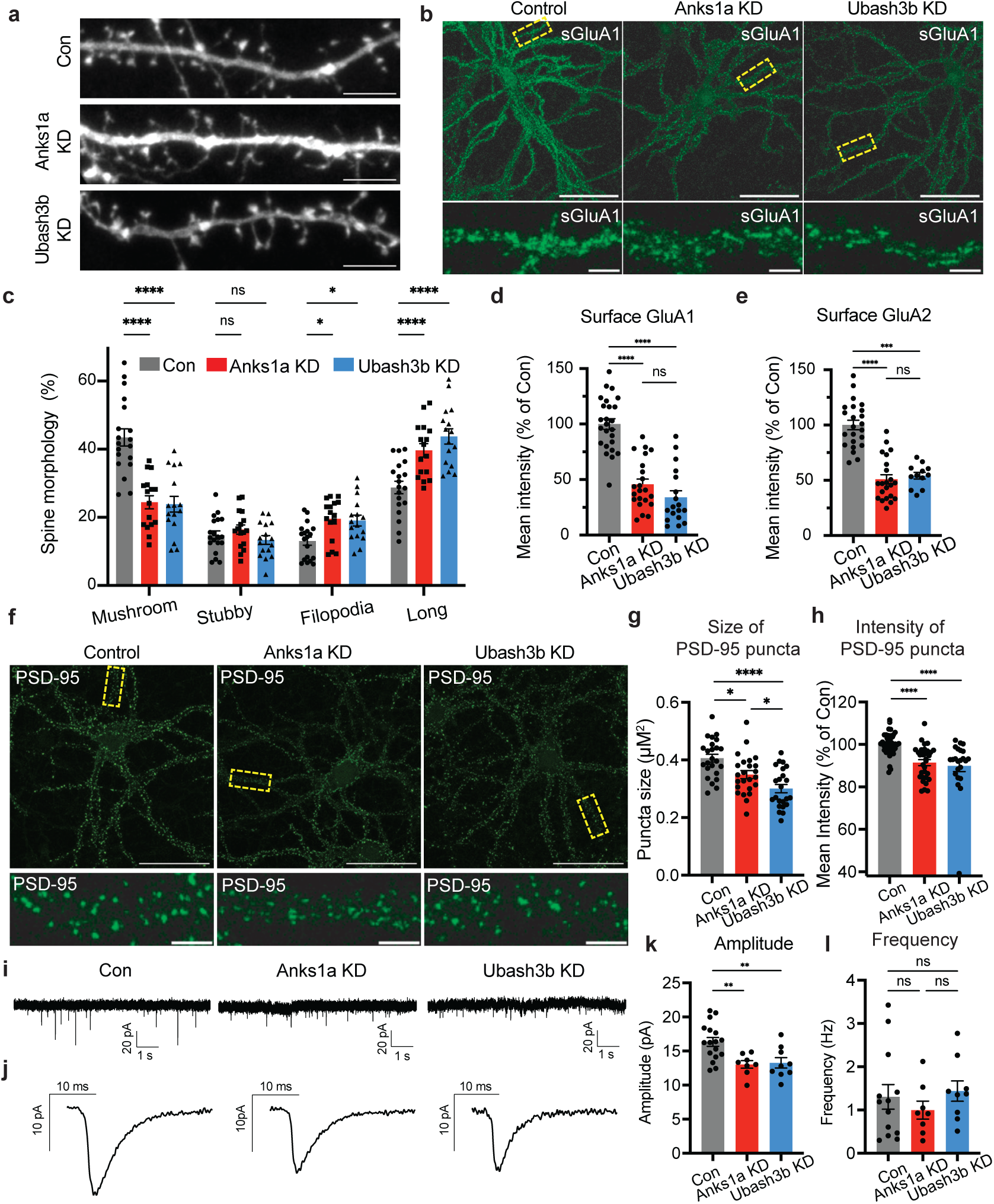
Loss of Anks1a or Ubash3b impairs synaptic structure and function. a–l, Primary mouse neurons expressing dCas9 via lentiviral transduction and gRNA via sparse transfection (a– b, i–l) or lentiviral delivery (c–h). a, c, Representative images (a) and quantification (c) of dendritic spines of DIV21 neurons. c, Two-way ANOVA with Dunnett’s multiple-comparisons test, *n* = 16–19 cells, 3 culture batches. b, d, e, Representative images (b) and quantification (d, e) of immunostaining of surface GluA1 (b, d) or surface GluA2 (e) puncta in DIV21 neurons. d, One-way ANOVA with Dunnett’s multiple-comparisons test, *n* = 18–25 cells from 3–4 culture batches. e, One-way ANOVA with Dunnett’s multiple-comparisons test, *n* = 12– 23 cells from 2–4 culture batches. f–h, Representative images (f) and quantification (g, h) of immunostaining of PSD-95 puncta in DIV21 neurons. g, h, One-way ANOVA with Dunnett’s multiple-comparisons test, *n* = 23–41 cells from 2–3 culture batches. i–l, Representative traces (i), representative mEPSC waveforms from one cell(j) and quantification of mEPSC amplitude (k) or frequency (l) from voltage-clamp recordings of DIV14–21 primary mouse neurons. k, l, One-way ANOVA with Dunnett’s multiple-comparisons test, *n* = 8–17 cells, 3–4 culture batches. Scale bars 50 μm (whole cell), 10μm (a) or 5 μm (dendrite zoom). Data are presented as mean ± s.e.m. ns, not significant. *P < 0.05, ***P < 0.001, ****P < 0.0001.

### Anks1a and Ubash3b promote activity-dependent accumulation of EGFR in synapses

In other cell types, Anks1a and Ubash3b enhance EGFR signaling by either enhancing EGFR recycling or preventing its endocytosis and degradation^26,27,35^. EGFR is also localized to synapses in neurons^8^, and pharmacological studies have shown that exogenous EGF facilitates LTP^40,41^, while EGFR inhibition impairs LTP^42^. These findings suggest that EGFR plays an important, yet largely overlooked, role in activity-dependent synaptic remodeling. We therefore hypothesized that Anks1a and Ubash3b enhances EGFR accumulation and signaling at synapses. To test this, we performed colocalization analyses with EGFR, as spatial proximity can indicate potential functional regulation. Since synaptic Anks1a and Ubash3b correlate with synaptic activity, we conducted immunostaining under baseline conditions to capture a range of activity states. Although most neurons showed sparse synaptic Anks1a and Ubash3b, we selected neurons with higher synaptic localization to enable more reliable colocalization analysis. Under these conditions, we observed robust colocalization of Anks1a with EGFR (Extended data Fig. 8a–b), and of Ubash3b with EGFR (Extended data Fig. 8c–d), suggesting that EGFR localization may be regulated by Anks1a and Ubash3b, and that EGFR is preferentially enriched at active synapses.

To test this directly, we treated cultured neurons with glutamate or cLTP and observed increased EGFR levels at synaptic regions (Fig. 5a–b and Extended Data Fig. 8e, g). Consistent with this, cLTP also increased synaptic phospho-Y1173 EGFR (Fig. 5c–d), an autophosphorylation site critical for downstream EGFR signaling^43^, indicating that EGFR is not only recruited to synapses but also locally activated in response to activity. Because EGF is one of the known ligands for EGFR, we also examined its synaptic distribution and observed increased EGF enrichment at synapses following activity (Extended Data Fig. 8f, h), supporting coordinated ligand-receptor regulation at active synapses.

**Figure 5.**
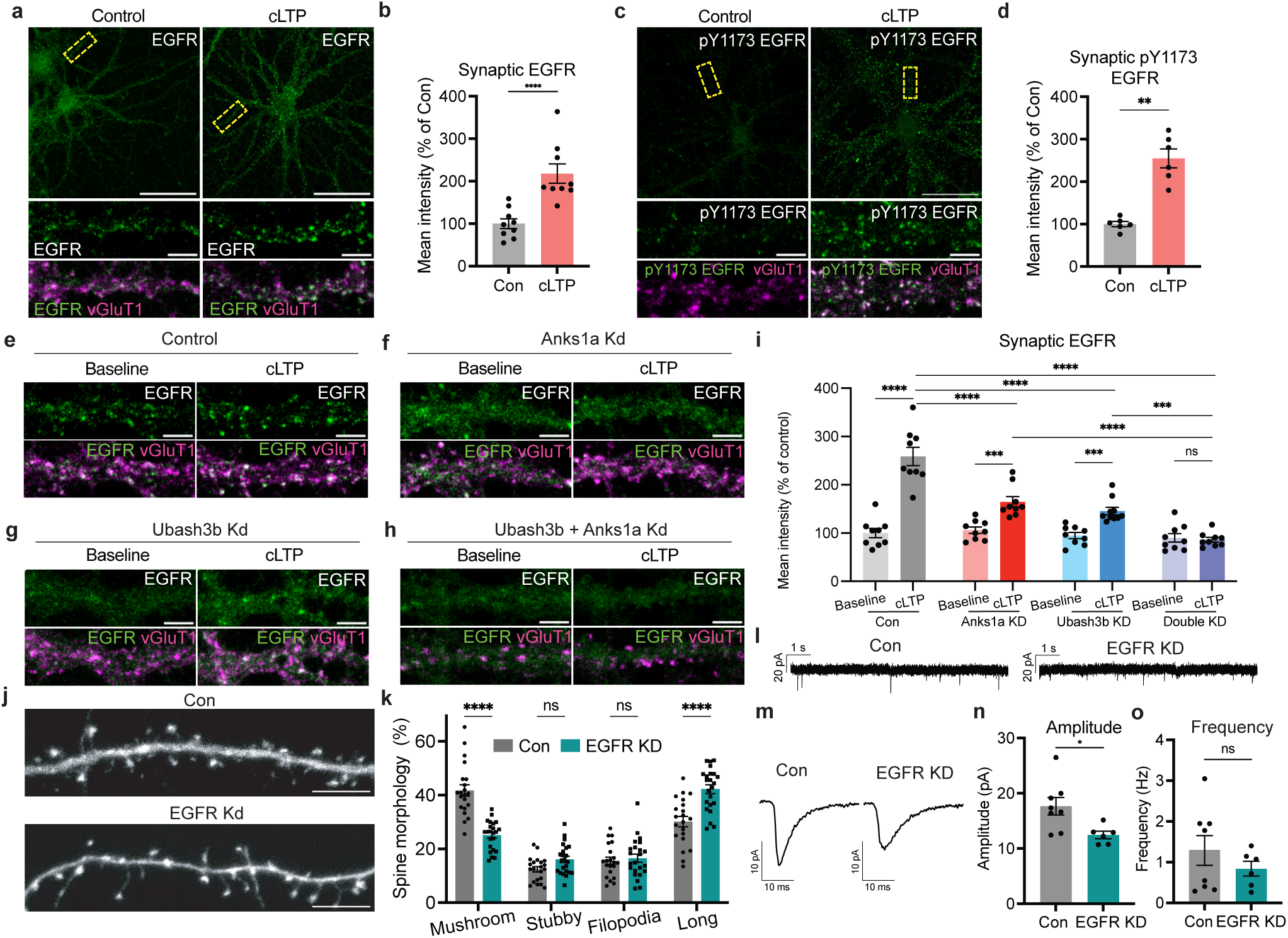
Activity-dependent EGFR accumulation at synapses requires Anks1a and Ubash3b. a–d, Representative images (a, c) and quantifications (b, d) of immunostained primary mouse neurons after chemical LTP (cLTP). b, Unpaired Welch’s t-test, *n* = 9, 3 culture batches. d, Unpaired Welch’s t-test, *n* = 6, 2 culture batches. e–i, Representative images (e–h) and quantification (i) of immunostained primary mouse neurons co-transduced with one lentivirus expressing dCas9 and another lentivirus expressing gRNA and subject to cLTP. l, Two-way ANOVA with Sidak’s multiple-comparisons test, *n* = 9–10 cells, 3 culture batches. j– k, Representative images (j) and quantification (k) of dendritic spines of mouse neurons transduced with lentivirus expressing dCas9 and transfected with gRNA construct. k, Two-Way ANOVA, *n* = 21–23, 3 culture batches. l–o, Representative traces (l), representative mEPSC waveforms from one cell (m) and quantification of mEPSC amplitude (n) or frequency (o) from voltage-clamp recordings of DIV13–14 primary mouse neurons. n, o, Mann-Whitney test, *n* = 6–8 cells, 2 culture batches. DIV21 primary mouse cortical neurons. Synaptic EGFR or pY1173 EGFR is defined as the signal within vGluT1 puncta. Scale bars 50μm (whole cell), 10μm (j) or 5μm (zoom and c, e–h). Data are presented as mean ± s.e.m. ns, not significant. **P < 0.01, ***P < 0.001, ****P < 0.0001.

To test whether Anks1a and Ubash3b are crucial for activity-dependent EGFR synaptic localization, we performed cLTP on Anks1a-KD and Ubash3b-KD neurons. Activity-induced EGFR enrichment at synapses was reduced but not abolished in either Anks1a-KD or Ubash3b-KD neurons (Fig. 5e–g, i and Extended Data Fig. 9a–c). We therefore tested whether these proteins act cooperatively by performing a double knockdown. This eliminated activity-induced EGFR enrichment at synapses (Fig. 5h–i and Extended Data Fig. 9d), indicating that Anks1a and Ubash3b function together in a partially redundant manner to jointly promote EGFR enrichment at synapses.

Finally, to determine whether EGFR is similarly required for synaptic maturation, we performed EGFR knockdown (Extended Data Fig. 9e–f). EGFR-KD neurons showed reduced mushroom spine abundance and increased long spines (Fig. 5j–k), together with decreased surface GluA1 and GluA2 levels (Extended Data Fig. 9h–k), indicative of impaired structural and functional synaptic maturation. mEPSC recordings further revealed reduced amplitudes without changes in frequency (Fig. 5l–o), consistent with weakened synaptic function without altered synapse number. Overall, these results reveal the importance of EGFR signaling in the maturation and functional strengthening of excitatory synapses.

### EGFR signaling is necessary for synaptic plasticity and learning and memory

Since synaptic EGFR signaling is rapidly induced by glutamate stimulation and cLTP (Fig. 5a–b and Extended Fig. 8e, g), we next asked whether EGFR signaling is required for synaptic plasticity. To avoid potential developmental effects of EGFR knockdown, we acutely inhibited EGFR using the selective inhibitor AG-1478. While cLTP increased surface GluA1 and GluA2 in control neurons, indicating increased synaptic strength, these increases were abolished by AG-1478 (Fig. 6a–c and Extended Fig. 10a–b), consistent with previous reports that EGFR activity is required for LTP^42^.

**Figure 6.**
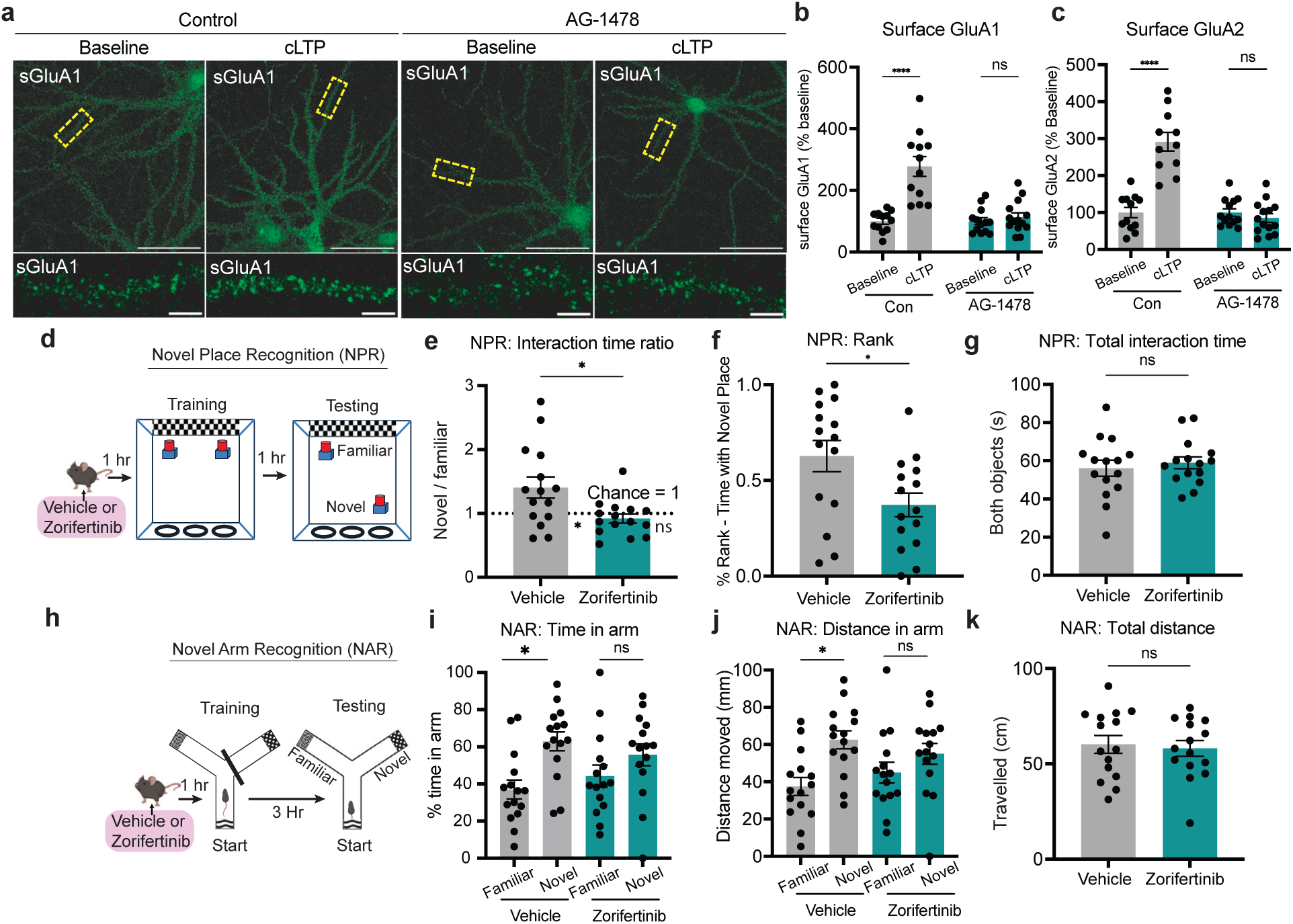
Acute EGFR inhibition impairs synaptic plasticity and learning and memory. a–c, Representative images (a) and quantifications (b, c) of immunostained primary mouse neurons treated with AG-1478 or control and immunostained for surface GluA1 (a–b) or surface GluA2 (c) 20 min after cLTP treatment. b, c, Two-way ANOVA, *n* = 11-14, 4 culture batches. d–k, 4–5-month-old mice treated with vehicle or zorifertinib were subject to novel place recognition (d–g) and novel arm recognition (h–k) tasks, *n* = 15 mice for each group. d, Experimental timeline for novel place recognition. e, Ratio of time interacting with the object in the novel versus familiar location. Welch’s t-test was used to compare groups. Dashed line indicates no preference (ratio = 1). One-sample t-test results comparing each group to 1 are shown below the dashed line. f, All mice were percentile-ranked based on percentage of time spent exploring the object in the novel location and scaled from 0–1, with higher values indicating better performance, Welch’s t-test. g, Total time mice spent interacting with both objects during testing. Welch’s t-test. h, Experimental timeline for novel arm recognition. i, Percentage of time in each arm, paired t-test. j, Distance moved in each arm, paired t-test. k, Total distance moved, Welch’s t-test. Scale bars 50μm (whole cell) or 5 μm (zoom). Data are presented as mean ± s.e.m. ns, not significant. *P < 0.05, ***P < 0.001, ****P < 0.0001.

Because LTP is widely considered a cellular substrate for learning and memory, we examined whether EGFR activity also contributes to learning and memory in vivo. To test this, we treated mice with zorifertinib (AZD3759), an EGFR inhibitor that permeates the blood-brain-barrier^44^, and assessed their performance in the novel place recognition (NPR) and novel arm recognition (NAR) tasks. Zoifertinib impaired recognition of the relocated object in NPR (Fig. 6d–f and Extended Fig. 10b–d) and disrupted the preference for the novel arm in NAR (Fig. 6h–j and Extended Fig. 10g–h), indicating deficits in spatial recognition memory. Importantly, zorifertinib did not affect locomotion, exploratory behavior, or anxiety-like behavior as measured in NPR and open field tests (Fig. 6g, k and Extended Figure 10e, f, i–m). Together, these findings reveal that EGFR signaling is crucial for synaptic plasticity and learning and memory.

## Discussion

Our study introduces synaptic Cal-ID as a strategy for activity-resolved proteomic characterization of active synapses under physiological conditions. Using this approach, we mapped the molecular landscape of active synapses in vitro and in vivo and uncovered an activity-dependent spatial regulation of EGFR signaling through the local enrichment of EGFR and its regulatory proteins, Anks1a and Ubash3b, that is required for synaptic maturation, plasticity and memory. These findings reveal a previously unrecognized mechanism by which synaptic activity establishes local growth factor signaling competence to regulate synapse function and behavior.

By labeling proteins during periods of synaptic activity, synaptic Cal-ID captures calcium-dependent molecular events with spatiotemporal precision, enabling the identification of molecular processes driving activity-dependent synaptic remodeling despite the spatially sparse and temporally transient nature of activity within intact circuits. Notably, labeling is initiated by exogenous biotin administration, which substantially increases labeling above the low basal signal generated by endogenous biotin. Our findings demonstrate how unbiased proteomic profiling of active synapses can uncover proteins and molecular pathways that are selectively engaged during activity to remodel synapses. Applying synaptic Cal-ID to specific neuronal populations or brain regions should further reveal the molecular mechanisms underlying the diverse forms of synaptic plasticity exhibited across neural circuits, an important challenge given the remarkable molecular diversity of synapses ^45,46^ and the distinct plasticity rules that govern them^46,47^.

Previous work has implicated synaptic EGFR signaling in plasticity, with EGFR detected at synapses^8^ and pharmacological modulation of its signaling influencing LTP^40–42^. However, how this broadly acting growth factor pathway is selectively engaged at active synapses has remained poorly understood. Through activity-resolved proteomics, we identified Anks1a and Ubash3b as previously unrecognized synaptic proteins that undergo rapid, NMDAR-dependent, and protein synthesis-independent enrichment following stimulation, revealing local recruitment of signaling proteins as an early molecular event in synaptic remodeling. Consistent with their established roles in regulating EGFR in other cellular contexts^26,27,35^, we found that Anks1a and Ubash3b promote activity-dependent synaptic enrichment and signaling of EGFR. We also observed an activity-dependent increase in synaptic EGF, suggesting that local ligand availability may further reinforce this spatial restriction, although the cellular source of EGF remains to be determined. Together, these findings support a model in which synaptic activity locally enriches EGFR, its regulatory machinery, and ligand to establish EGFR signaling competence at active synapses. More broadly, active synapses can confer spatial specificity on a broadly expressed growth factor pathway by locally concentrating the molecular components required for its activation.

Our results further suggest that physiological EGFR signaling contributes to synaptic plasticity and cognitive function in the adult brain. Acute EGFR inhibition impaired both cLTP-induced AMPAR recruitment and spatial memory, extending previous studies linking EGFR signaling to LTP^42^ and learning in *Drosophila*^48^. These findings may also have implications for EGFR-targeted cancer therapies. EGFR is a well-established oncogenic driver, and many EGFR-driven cancers harbor activating mutations that can be selectively targeted. The development of next-generation EGFR inhibitors with high blood–brain barrier permeability, such as zorifertinib, has improved the treatment of central nervous system (CNS) metastases^49^ by achieving therapeutic drug concentrations within the brain. Our findings raise the possibility that increased inhibition of physiological EGFR signaling in the brain could affect synaptic and cognitive function. Thus, as brain-penetrant EGFR therapies continue to advance, our findings highlight the potential importance of selectively targeting oncogenic EGFR while preserving physiological EGFR signaling in the brain. Mutant-selective, brain-penetrant inhibitors such as osimertinib^50^, illustrate the feasibility of this therapeutic strategy.

Although depletion of Anks1a or Ubash3b did not completely abolish activity-induced EGFR enrichment, depletion of either protein markedly impaired synaptic maturation and function. One possibility is that partial disruption of activity-dependent EGFR signaling is sufficient to impair normal synaptic development and plasticity. Alternatively, Anks1a and Ubash3b may regulate additional EGFR-independent mechanisms that contribute to synaptic function. Distinguishing between these possibilities will require further investigation.

In summary, this study uses calcium-dependent proximity labelling to examine the molecular architecture of active synapses and uncovers a previously unrecognized activity-dependent EGFR signaling pathway involving Anks1a and Ubash3b. More broadly, our findings reveal a previously unappreciated layer of synaptic regulation in which activity rapidly recruits local signaling machinery to active synapses to promote synaptic remodeling, plasticity, and memory.

### Limitations of study

Three considerations are important when applying synaptic Cal-ID. First, its calcium-responsive domains are derived from GCaMP6s^51^, thus its calcium sensitivity is expected to fall within a similar range, although the precise affinity has not been directly measured. Synaptic Cal-ID nevertheless responds to physiologically relevant calcium elevations within synapses, including those induced during LTP, and its labelling efficiency is likely influenced by the magnitude of local calcium signals. Second, synaptic Cal-ID labeling is cumulative over the labeling period and therefore reflects integrated rather than instantaneous synaptic activity.

Synapses experiencing more frequent or sustained calcium elevations are expected to accumulate greater labeling, such that protein recovery may be influenced by both the number and magnitude of local calcium events. Thus, synaptic Cal-ID preferentially captures the molecular composition of synapses with greater cumulative activity during the labeling window rather than providing a binary measure of synapse activation. Lastly, because TurboID labeling is proximity-dependent with nanometer-scale spatial resolution^52^, the proteins captured by synaptic Cal-ID are determined in part by its subcellular localization. In this study, targeting Cal-ID to PSD-95 likely biases detection toward proteins within or proximal to PSD-95–associated synaptic nanodomains. Alternative targeting strategies could therefore extend synaptic Cal-ID to resolve activity-dependent protein composition across distinct synaptic nanodomains.

## Methods

### Primary neuron culture

Cortices with hippocampi were dissected from E17 CD1 mouse embryos (Charles River) in dissection medium HBSS containing Pen-strep, 1mM sodium pyruvate, 20mM HEPES, 25mM glucose and digested with 20 units/ml papain (Worthington) for 15 min at 37°C. Neurons were plated at a concentration of 32,000–80,000 neurons/cm^2^ with higher densities for biochemistry and lower densities for immunostaining. Culture plates and coverslips were coated with 0.5 mg/mL poly-D-lysine dissolved in Trizma buffer. Cultures were maintained in Neurobasal Plus medium with B-27 plus and 0.25x GlutaMax. Neurons were treated on DIV5 with 1.67µM FdU (5-Fluro-20-deoxyuridine and Uridine) to stop glia proliferation and fed twice weekly.

### Animals

All animal housing and handling procedures were conducted according to protocols approved by the Institutional Animal Care and Use Committee (IACUC) at the University of California, San Francisco. C57BL/6J mice were group housed (2–5 per cage) and given water and food ad libitum on a 12/12 h light/dark cycle in a temperature-controlled (22–24 °C) and humidity-controlled (40–60%) environment. During breeding for P0 pup injections, mice were provided with forage mix, hut and Love Mash (Bio-Serv).

### Cloning and plasmids

Polymerase chain reactions were performed with Q5 polymerase (NEB) and the fragments were assembled using In-fusion (Takara). The ligated plasmid products were introduced by heat shock transformation into competent NEB Stable cells (NEB). gRNAs cloned into lenti U6-sgRNA/EF1a-mCherry (Addgene 114199). dCas9 cloned from pLX303-ZIM3-KRAB-dCas9 (Addgene 154472).

### Lentivirus generation

HEK293FT cells were maintained in DMEM with L-Glutamine, 4.5g/L Glucose and Sodium Pyruvate with GlutaMax, Pen-Strep, and 10% FBS and passaged 2–3x a week. For lentivirus production, cells were cultured on 1% gelatin-coated dishes in medium containing 25 mM HEPES without pen/strep. Transfer vectors were co-transfected with second-generation VSV-G and gag/pol packaging plasmids (1:1:1.2 molar ratio) using JetOPTIMUS (Polyplus). Viral supernatants were collected 48 and 72 h after transfection, filtered, concentrated using Lenti-X Concentrator (Takara), resuspended in Neurobasal Plus medium, and titrated using Lenti-X GoStix (Takara).

### Neuron transfection

Neurons were transfected between DIV 1–5 using Lipofectamine 2000. Lipofectamine and DNA were separately diluted in Neurobasal media (no B27) then combined and incubated at 37 °C for 20 min. Some media from each well containing neurons collected, mixed with fresh media and retained. Complexes were added dropwise to neurons and incubated for 2 h. Cells were washed once with fresh media and returned to the conditioned(old)/fresh media mixture.

### Intracranial neonatal mouse injections

Neonatal mice (P0–1) were cryo-anesthetized by placing on ice for ∼5 min. When the animal was fully anesthetized, confirmed by absence of pedal withdrawal reflex, it was gently placed in a head mold on top a cold aluminum plate. Each pup received a total of 1.4µL of AAV9 (HHMI Viral Tools, Cal-ID-PSD-95: 2.06×10^13^ GC/mL, cpTurboID-PSD-95: 1.81×10^13^) in bilateral injections, delivered as four 350-nL injections at a rate of 250 nL/minute. Two injections were made into each cortex, located at one-third and two-thirds the distance from bregma to lambda. approximately 0.6 mm lateral to the sagittal suture and 0.6 mm ventral to the skull surface.

### Biotin injection and enriched environment

USP-grade biotin was dissolved in saline and sodium hydroxide, adjusted to pH 7.5, and administered intraperitoneally at 24 mg/kg. Mice (2–5 per cage) were transferred to enriched environment cages containing bedding from the home cage, forage mix, food, hydration gel, and a variety of enrichment objects (Bio-Serv) that promoted running, exploration, gnawing, sheltering, and vertical climbing (Extended Data Fig. 3e).

### Perfusion and immunohistochemistry

Mice were deeply anesthetized with isoflurane and transcardially perfused with ice-cold PBS followed by 4% paraformaldehyde (PFA)/4% sucrose in PBS. Brains were post-fixed overnight at 4 °C, cryoprotected in 30% sucrose, and sectioned sagittally at 40 μm on a freezing microtome (Leica CM3050S). Brain sections were mounted onto slides and air-dried prior to immunostaining. Mounted sections were incubated with primary antibodies overnight at 4 °C, followed by secondary antibodies for 2 h at room temperature, and coverslipped with mounting medium containing DAPI.

### Subcellular fractionation

All procedures were performed on ice or at 4 °C in the presence of phosphatase and protease inhibitors, 5 mM sodium pyrophosphate, 1 mM EDTA, and 1 mM EGTA. Cultured cortical neurons and mouse brains were homogenized in 320 mM sucrose, 10 mM HEPES using a 26-gauge needle or glass homogenizer, respectively. Homogenates were centrifuged at 800 × g for 10 min to generate the post-nuclear pellet (P1) and supernatant (S1). S1 was centrifuged at 15,000 × g for 20 min to yield the crude membrane fraction (P2) and cytosolic fraction (S2).

For cultured neurons, P2 was osmotically lysed and centrifuged at 25,000 × g for 20 min to generate LP1 (mitochondria, pre- and postsynaptic membranes) and LS2 (synaptosomal cytosolic fraction). For mouse brains, P2 was purified by discontinuous sucrose density-gradient ultracentrifugation (0.8, 1.0, and 1.2 M sucrose) at 82,500 × g for 2 h, and synaptosomes collected from the 1.0/1.2 M interface were pelleted at 100,000 × g for 30 min to generate LP1. LP1 fractions were extracted with 0.5% Triton X-100 for 5 min (cultured neurons) or 10 min (brain) and centrifuged at 25,000 × g for 20 min to separate presynaptic membranes (supernatant) from the postsynaptic density (pellet).

### Sample preparation for quantitative mass spectrometry

#### Samples for mass spectrometry were prepared in biological triplicates

Cultured neurons: 2 × 150 mm dishes of neurons per sample. Lentivirus containing Cal-ID-PSD95 or cpTurboID-PSD95 were added to neuronal cultures on DIV 7. Biotinylation (100 µM biotin, 1 hour in media) and lysis on DIV 21.

Mouse brains: 6 male 4–5-month-old C57BL/6J mice per sample. Biotin administered intraperitoneally and immediately placed in the enriched environment cage for 4 hours. Mice brains were harvested, cortices dissected and immediately frozen in ethanol/dry ice and stored in −80 °C. Subcellular fractionation was performed from thawed cortices.

Following subcellular fractionation, PSD pellet was resuspended in 0.2% sodium laurate lysis buffer (50 mM NaF, 5 mM NaPPi, 1 mM EDTA, 1 mM EGTA in PBS pH 7.2). The solution was sonicated, clarified by centrifugation (20,000 × g, 10 min), and protein concentrations determined by BCA assay. Biotinylated proteins were enriched using streptavidin magnetic beads (Pierce), and enrichment efficiency was verified by western blot. Beads were washed sequentially with lysis buffer, 1 M KCl, 0.1 M Na₂CO₃, 2 M urea, lysis buffer, 50mM Tris-HCl, and water.

#### On beads digestion, TMT labelling and on-tip high pH C18 chromatographic fractionation

Samples were prepared similar to the published paper^53^ with the following modifications. Beads resuspended in 15 µl TCEP, second on bead digest with 9 µl 20 mM TEAB and labelling reactions were combined and diluted with 1600 µl formic acid. Desalting: the OMIX tip was wet with 0.1% formic acid in 95% MeCN, then equilibrated in 0.1% formic acid as indicated by the manufacturer. Sample was loaded in the tip resin by passing it through it 4 times. The tip was then washed 3 times with 200 μL 0.1% formic acid. Then, the tip was washed with 2 aliquots of 40 ul 20 mM ammonium formiate. These aliquots were collected and kept for MS analysis.

Peptides were then eluted sequentially in 20 mM ammonium formiate solutions with increasing amounts of MeCN (2.5%, 5%, 7.5%, 10%, 12.5%, 15%, 17.5%, 20%, 90%), in 2 x 40 μl consecutive aliquots for each collected in the same tube, and the resulting 10 fractions dried and resuspended in 2.5 μl 0.1% formic acid for mass spectrometry analysis.

### Mass spectrometry and data analysis

Samples were processed similar to the previous study^53^ with modifications. Samples run in either Orbitrap Exploris 480 (Thermo Scientific) or a Orbitrap Lumos Fusion (Thermo Scientific). MS/MS spectra were acquired in centroid mode with resolution 60000 from m/z=120 in the Exploris, 50000 from m/z=110 in the Lumos. Peak lists were generated using PAVA in-house software. All generated peak lists were searched against the mouse subset of the SwissProt database (SwissProt.2019.07.31), using Protein Prospector with the following parameters: Enzyme specificity was set as Trypsin, and up to 2 missed cleavages per peptide were allowed.

Carbamidomethylation of cysteine residues, and TMTPro16plex labeling of lysine residues and N-terminus of the protein were allowed as fixed modifications. N-acetylation of the N-terminus of the protein, loss of protein N-terminal methionine, pyroglutamate formation from of peptide N-terminal glutamines, and oxidation of methionine were allowed as variable modifications Mass tolerance was 5 ppm in MS and 30 ppm in MS/MS. The false discovery rate was estimated by searching the data using a concatenated database which contains the original SwissProt database, as well as a version of each original entry where the sequence has been randomized. A 1% FDR was permitted at the protein and peptide level. For quantitation only unique peptides were considered. Relative quantization of peptide abundance was performed via calculation of the intensity of reporter ions corresponding to the different TMT labels, present in MS/MS spectra. Intensities were determined by Protein Prospector. Summed intensity per sample on each TMT channel for all identified spectra belonging to endogenously biotinylated proteins (carboxylases) were used to normalize individual intensity values. Relative abundances were calculated as ratios relative to the no virus samples. Spectra representing replicate measurements of the same peptide were kept. Individual spectral ratios were aggregated to the peptide level using median values of the log_2_ ratios, and these values were in turn used to calculate the protein levels log_2_ ratios (as medians of the individual peptide levels of each protein).

For differential expression analysis, contaminants and proteins that had <2 unique peptide were filtered. For each Syn-Cal-ID and Syn-cpTurboID sample, protein enrichment levels were normalized to the sample median, and the median enrichment value across triplicates was used for each protein. Loess regression (α = 0.3) of Syn-Cal-ID versus Syn-cpTurboID enrichment was performed and the difference between observed Syn-Cal-ID enrichment for a protein and the loess prediction based on its Syn-cpTurboID enrichment was used to quantify relative Cal-ID enrichment.

### STRING analysis

Protein–protein interaction networks and functional enrichment analysis for Gene Ontology (GO) biological processes were performed using the STRING database via the Cytoscape stringApp (v2.2.0) with a whole-genome background. Enrichment significance is reported as false discovery rate (FDR) provided by STRING. Networks were visualized in Cytoscape.

### Immunoblotting

Lysis in RIPA buffer with 1 % SDS (20 mM Tris-HCl (pH 7.4), 150 mM NaCl, 1 mM EDTA, 1 % NP-40, 1 % sodium deoxycholate, 1 % SDS, protease inhibitors). Lysates were incubated on a rotator for 15 min 4 °C, supernatant collect after centrifugation for 10 min × 17,000 xg at 4 °C. protein concentration was measured, and the lysate was mixed with 4 × NuPAGE LBS sample buffer with DTT. Samples were separated using SDS-PAGE on Invitrogen Bolt™ system with Bis-Tris 4–12% gradient gels and transferred onto polyvinylidene difluoride (PVDF) membranes using a Bio-rad transblot electrophoretic transfer system. Membranes were blocked in Intercept® TBS Blocking Buffer (Licor). Li-Cor Odyssey Clx imager was used for visualization.

### RT-PCR

Neurons were lysed with CellAmp Direct Prep Kit for RT-PCR & Protein Analysis (Takara) followed by gDNA elimination and reverse transcription using PrimeScript™ RT Reagent Kit with gDNA Eraser (Takara). qPCR primers were designed using Primer-BLAST (NCBI). qPCR performed using PowerUP SYBR Green Master Mix (Thermo Fisher A25742)) using Applied Biosystems QuantStudio7 system. Relative expressions of genes were calculated using the delta-delta Ct method with the GAPDH as the normalizing gene.

### Immunocytochemistry

Neurons were fixed in 4% PFA/4% sucrose 10–15 min (other antibodies) or 4% glyoxal pH 4.5 (pY1173 EGFR staining) for 5 min. Cells were permeabilized in 0.25% Triton-X (all other antibodies) or 0.1% Triton-X (EGFR) and blocked for 1 hour in 5 % BSA/5% normal goat serum. Primary antibody incubation was overnight in 1% BSA at 4 °C, secondary antibodies in 1% BSA for 1 hour at room temperature.

For surface AMPAR and EGF staining, neurons were fixed in 4% PFA/4% sucrose for 4 min. EGF staining included permeabilization with 0.1% Triton X-100 for 15 min, whereas surface AMPAR staining was performed without permeabilization. Blocking in 5% BSA/5% normal goat serum for 30 min, before primary antibody incubation in 1% BSA/1% NGS for 2 h at room temperature or 4 °C overnight. Neurons were permeabilized with 0.25% Triton X-100 for 15 min followed by blocking as before and other primary antibody incubation overnight in 1% BSA at 4 °C. Secondary antibodies in 1% BSA for 1 hour at room temperature.

### Imaging and analysis

Images were taken with SP8 (Leica) confocal laser scanning microscope with a 20x, 40x or 63x oil objective. All image analysis was done using FIJI/ ImageJ software. Image acquisition and analysis parameters, including thresholding and ROI selection criteria, were kept constant across all conditions within each experiment.

Synaptic regions of interest (ROIs) were generated with Analyze Particles based on immunostaining of synaptic markers. PSD95 was used as the primary postsynaptic marker when signal quality was sufficient. In experiments involving multiplex immunostaining conditions that compromised PSD95 signal preservation, vGluT1 was used as an alternative synaptic marker to define puncta. For each image, a region within the dendrite without clear punctate signal was measured and used for background subtraction.

Surface AMPAR quantification was performed by manually tracing non-primary dendritic segments (≥5 per image) with clear surface staining. The Analyze Particles function was then applied to isolate puncta, and integrated density (mean fluorescence intensity × puncta area) was measured and normalized to dendrite length.

For PSD95 puncta quantification, Analyze Particle function was used and resulting ROIs were used as a mask to quantify to mean fluorescent intensity and area of the PSD95. Puncta density was calculated by total number of puncta divided by dendrite lengths.

Colocalization analysis was performed using JaCOP with manual thresholding. Costes randomization performed using xy block size 2, 10 rounds, pad with black pixels, last randomized image used for colocalization quantification control.

Spine morphology was analyzed manually from maximum intensity projected images by two blinded annotators. Spines were classified as mushroom, stubby, long/thin, or filopodia based on visual assessment of spine shape, guided by approximate morphological criteria that would correspond to the following measurements if quantified: mushroom (spine length < 3 µm and maximum head width > 2× mean neck width), stubby (spine length < 1 µm), long/thin (head width ≥ neck width), and filopodia (head-to-neck width ratio < 1.2).

### Intracellular whole-cell recordings

Experiments performed as in previous study^54^. Whole-cell recordings were obtained at room temperature from cortical neuronal cultures in ACSF solution (130mM NaCl, 3mM KCl, 10 mM HEPES, 10 mM D-glucose, 2 mM MgSO_4_, and 2 mM CaCl_2_, pH 7.4, 100 μM picrotoxin, 1 μM TTX, and 100 μM DL-APV). Glass patch pipettes with resistances of 3–6 MΩ were filled with (in mM) 115 CsMeSO_4_, 8 CsCl, 5 TEA-Cl, 0.5 EGTA, 10 HEPES, 1 QX-314, 10 Na phosphocreatine, 0.5 Na-GTP, and 4 Mg-ATP, pH 7.2, at 290 mOsm. mEPSCs were detected by template matching using Clampfit 10 (Molecular Devices). Kinetic measurements were performed on mean mEPSC traces. Tau was measured using a single exponential decay function. Mouse primary cortical neurons were cultured either in Neurobasal plus (DIV19–21) or NbActiv4 (Transnetyx; DIV12–14), which supports robust electrophysiological recordings. As electrophysiological properties were comparable between conditions, datasets were pooled for analysis.

### Drug treatments in culture

DIV 19–21 mouse primary cortical neurons were pre-equilibrated with artificial cerebrospinal fluid (ACSF) for 20 min at 37 °C. ACSF contained 143 mM NaCl, 5 mM KCl, 10 mM HEPES, 10 mM glucose, 2 mM CaCl₂ and 1 mM MgCl₂ (300–310 mOsm, pH 7.4). For inhibition experiments, drugs (Anisomycin, APV or TTX) were added during this period. For cLTP experiments: all media also contained 500 nM tetrodotoxin, 20 μM bicuculine, and 1 μM strychnine. After 20 min pre-equilibration, neurons were switched to Mg²⁺-free ACSF containing 200 μM glycine for 5 min to induce cLTP. For labeling, 100 μM biotin was added 5 min prior to cLTP or BAPTA-AM/drug cocktail treatment, followed by incubation to yield total labeling times of 30 min (cLTP) or 60 min (BAPTA-AM/drug cocktail).

### Mouse behavioral assays

Mice were administered vehicle (20% SBE-β-CD in sterile saline) or zorifertinib (15 mg/kg) by oral gavage once daily at a dosing volume of 6 mL/kg 1 hour prior to the first training/testing for that day. Behavioral assessments were conducted 1–4 hours after dosing, a time window previously shown to provide effective drug exposure^44^. For all tests, mice were acclimated to the testing room for 1 h before.

Open-field: acrylic chamber (43 × 43 × 30 cm) equipped with photobeam arrays to detect horizontal and vertical movement. Mice were allowed to freely explore the arena for 15 min. Locomotor activity was quantified by beam breaks and video tracking with Activity Monitor Software (Med Associates).

Novel Place Recognition: white acrylic arena (40 × 40 × 22 cm) containing two identical objects with identical odors and two spatial cues on opposite walls. Day 1, 15 min habituation. Day 2, 10-min training session. Day 3, 10 min training, a 10-min test session 1 hr later in which one object was relocated to a novel position. CleverSys TopScan was used to analyze and quantify the data. Interaction time was defined as time spent sniffing within a 1.5cm radius around the objects.

Novel arm recognition: white plastic Y-maze consisting of three identical arms (49.5 × 8.0 × 23.5 cm) extending from a central triangular area. Distinct visual cues were placed at the ends of the three arms. During training (5 mins), one arm was blocked off. 3 h later, the arm was unblocked, and the mouse was allowed to freely explore all three arms for 5 min (test). Behavior was analyzed during the first 2 min of the test session because novelty preference is most sensitive immediately upon re-exposure to the maze. Noldus Ethovision was used to analyze and quantify the data.

All apparatus were cleaned with Vimoba before and after daily use and with 70% ethanol between animals.

### Quantification and statistical analysis

All data were presented as mean ± SEM (standard error of mean). Statistics were performed using GraphPad Prism. Statistical significance was defined as *, p<0.05; **, p<0.01, ***, p< 0.001, ****p<0.0001; n.s, no statistical significance. Unless data was shown to be normally distributed and have equal variances, non-parametric tests were used. For experiments with more than two groups, one-way ANOVA or two-way ANOVA was used.

## Resource Availability

### Lead contact

Requests for further information and resources should be directed to and will be fulfilled by the lead contact, Adeline J.H. Yong.

### Materials availability

Plasmids generated in this study will be deposited with Addgene and made publicly available upon publication.

### Data and code availability

The raw proteomics data have been deposited into ProteomeXchange/PRIDE with identifier PXD079330. All other raw data and analyzed data are available upon request.

## Acknowledgements

We appreciate all members of the Jan laboratory for their support and feedback, and Marena Tynan-La Fontaine, Armando Martinez, and Xingnu (Jessie) Zhai for administrative and laboratory support. We are grateful to Graeme W. Davis for his guidance and thoughtful discussions, and Franck Polleux, Peter H. Chipman and Fangming Xie for critical reading of the manuscript. We thank Iris Lo from Gladstone Institutes Behavioral Core for interpretation of behavior experiments. This work was supported by NIH/NINDS grants to Y.N.J. (R35NS097227 and R35NS137312), A.J.H.Y. (K99NS140624), N.T.I. (R21NS112842), L.Y.J (R35NS122110 and R01NS116588), and UCSF Sandler PBBR Postdoc Independent Research Award (A.J.H.Y), Weill Neurohub Next Great Ideas Program (N.T.I and Y.N.J.), Brain & Behavior Research Foundation Young Investigator Grant (J.W.K.), Dr. Miriam and Sheldon G. Adelson Medical Research Foundation (A. L. B), and Howard Hughes Medical Institute (L.Y.J. and Y.N.J.).

## Author contributions

A.J.H.Y. and Y.N.J. conceived of the study. A.J.H.Y. performed all experiments other than those stated. Y.W. performed electrophysiology experiments, and A.J.H.Y., and Y.W. performed the analysis. J.A.O-P. performed mass spectrometry and J.A.O-P., J.W.K., and N.T.I. performed proteomic data analysis. A.J.H.Y. and T.C. performed molecular cloning. A.J.H.Y, C.C. and M.V. Z. performed imaging analysis. L.D. performed and analyzed behavior experiments, and A.J.H.Y. and J.S. designed and interpreted behavior experiments. Y.N.J., N.T.I., L.Y.J., A.L.B., provided resources and scientific guidance. J.W.K. contributed to project design, discussion and data interpretation. A.J.H.Y wrote the manuscript with input from all authors.

---

**Extended Data Figure 1.**
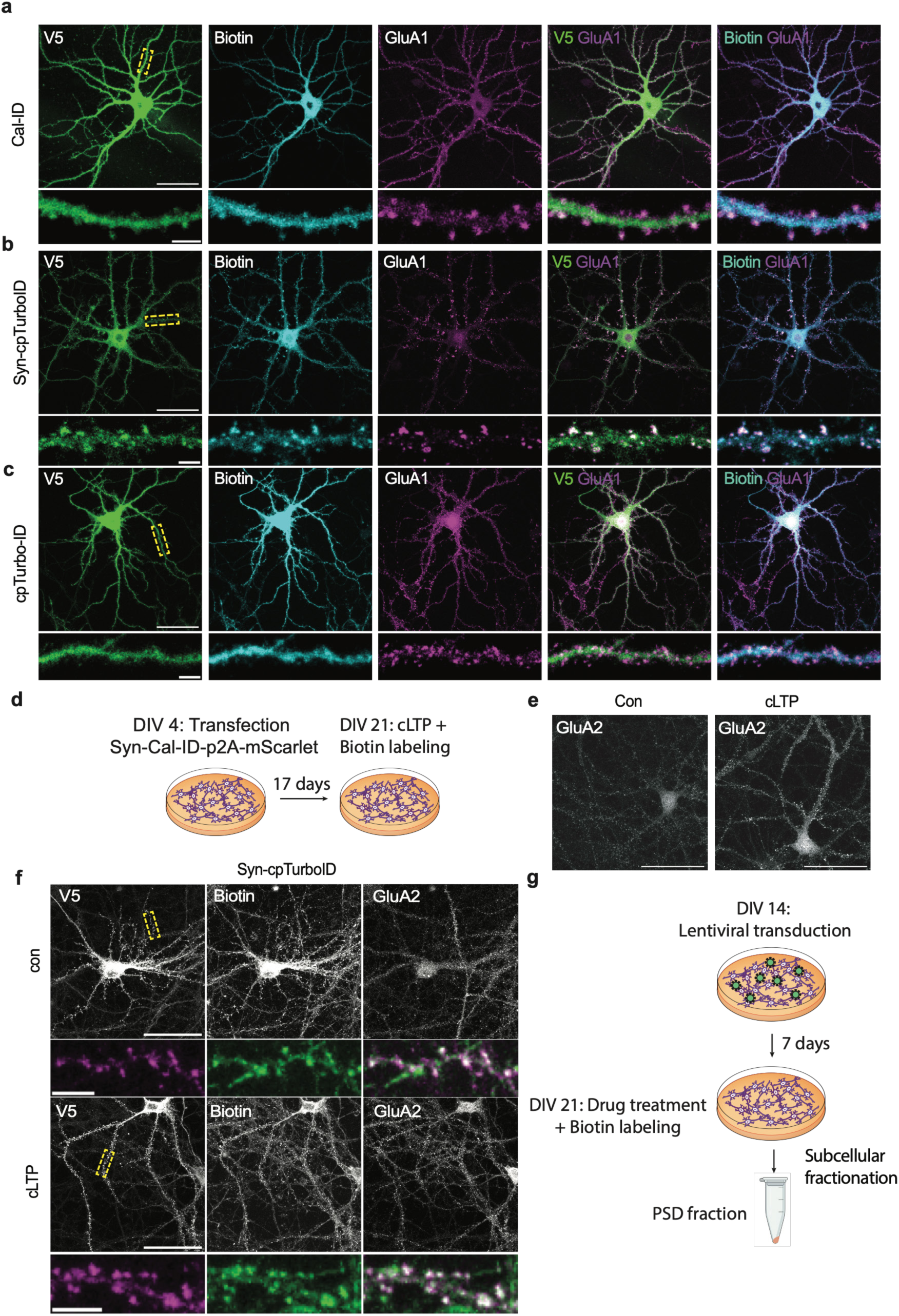
Synaptic Cal-ID is a synapse-localized calcium-dependent biotin ligase. a–c, Representative images of DIV21 immunostained mouse primary neurons transduced with virus expressing Cal-ID-V5 (a), Syn-cpTurboID-V5 (b) or cpTurboID-V5 (c). d, Experimental outline of cLTP experiments. e–f, Representative image of immunostained mouse neurons expressing Syn-Cal-ID (e) or Syn-cpTurboID (f). g, Experimental outline of experiments to test Syn-Cal-ID response to decreased calcium.

**Extended Data Figure 2.**
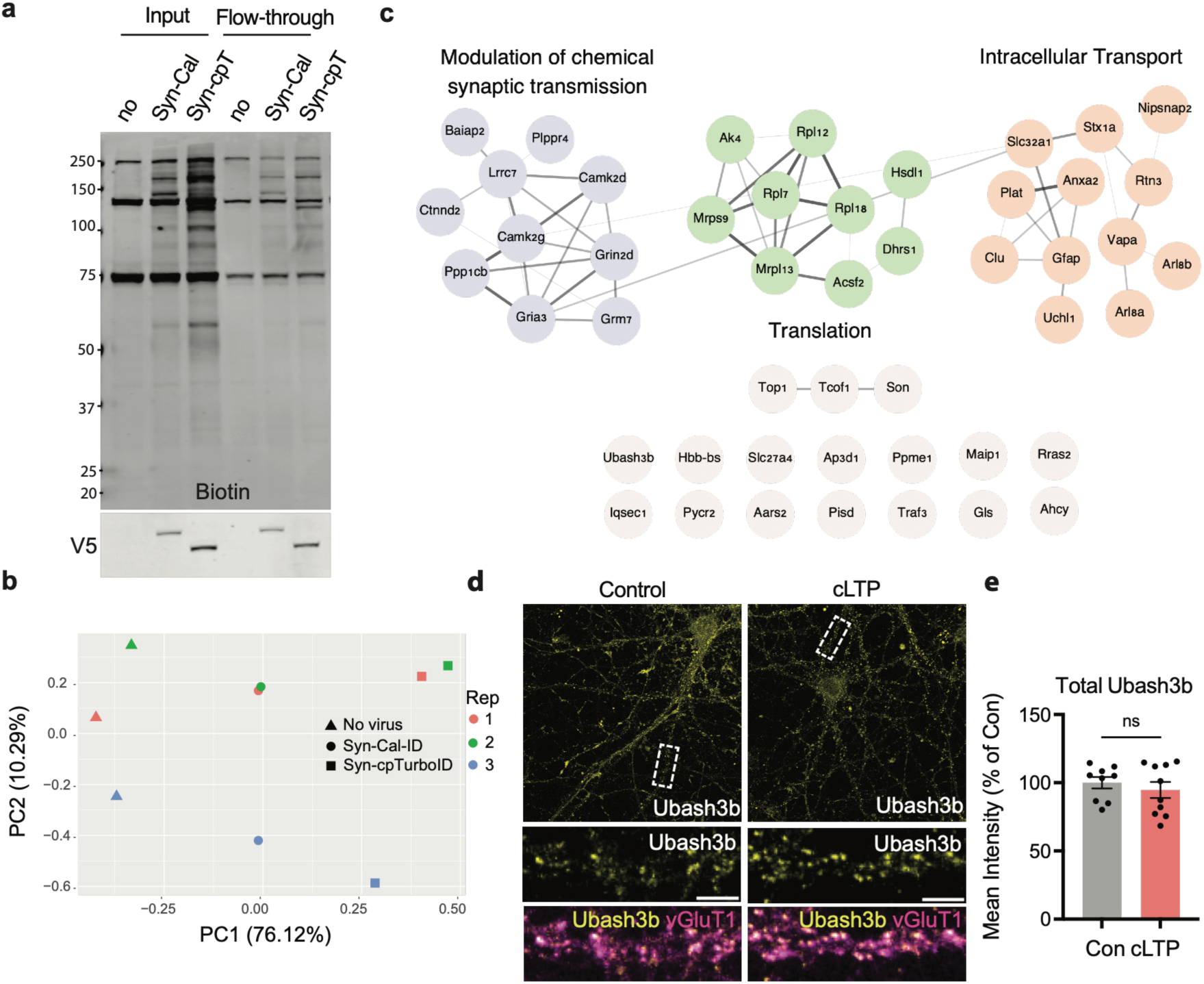
Proteomic screen using synaptic Cal-ID in vitro identifies Ubash3b as a novel activity-dependent synaptic protein. a, Immunoblot of PSD proteins from DIV21 mouse primary neurons pre- and post- streptavidin bead pulldown. B, Principal component analysis (PCA) of mass spectrometry samples. c, STRING network analysis of the top 4.7% of proteins ranked by enrichment ratio (confidence score cutoff = 0.29). Proteins cluster into Gene ontology (GO) biological processes including modulation of chemical synaptic transmission (FDR=0.0225), translation (FDR = 0.0035), intracellular transport (FDR = 0.0126). Node colors indicate functional categories; grey nodes represent unclustered proteins. Edges represent protein–protein associations, edge thickness and opacity scale with STRING confidence score. d, e, Representative image (d) and quantification (e) of DIV21 primary mouse neurons immunostained after cLTP. e, Mann-Whitney test, *n* = 9–10, 3 culture batches. Scale bars 50μm (whole cell) or 5 μm (zoom). Data are presented as mean ± s.e.m. ns, not significant.

**Extended Data Figure 3.**
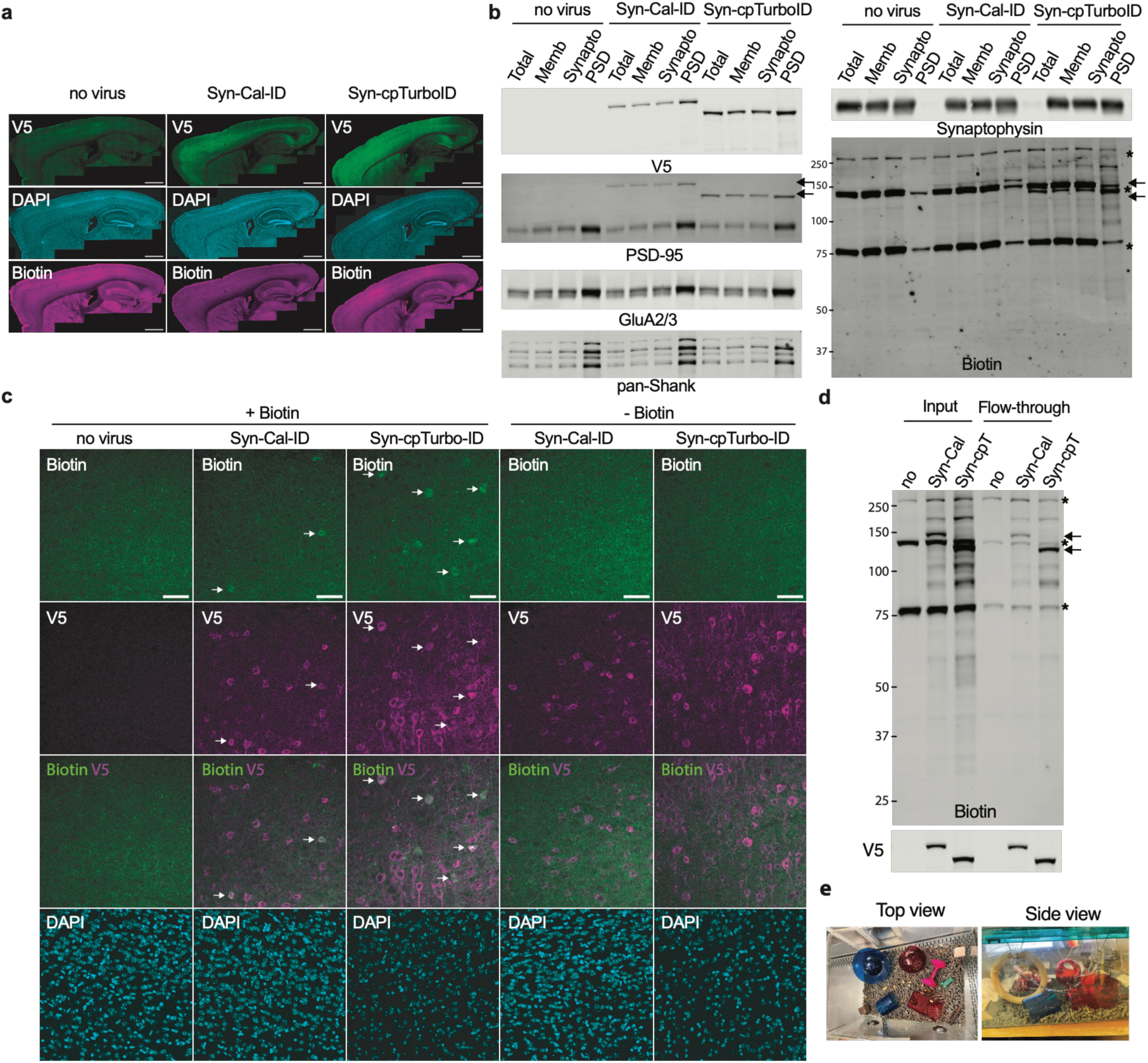
Validation of Syn-Cal-ID–mediated neuronal biotinylation in vivo. a,b, Representative images of brains of mice injected with Syn-Cal-ID or Syn-cpTurboID or no virus showing viral spread throughout cortex (a) and biotinylation of neurons (b). Due to background from endogenous biotinylated proteins, differences in labelling are most apparent at higher magnification, where regions of V5 signal overlap with biotinylation (white arrows). c, d, Immunoblots of subcellular fractions (c) or pre- and post-streptavidin bead pulldown from PSD fraction (d) from cortices of mice injected with Syn-Cal-ID or Syn-cpTurboID or no virus. e, Enriched environment cages. Scale bars 1mm (a), 50μm (b).

**Extended Data Figure 4.**
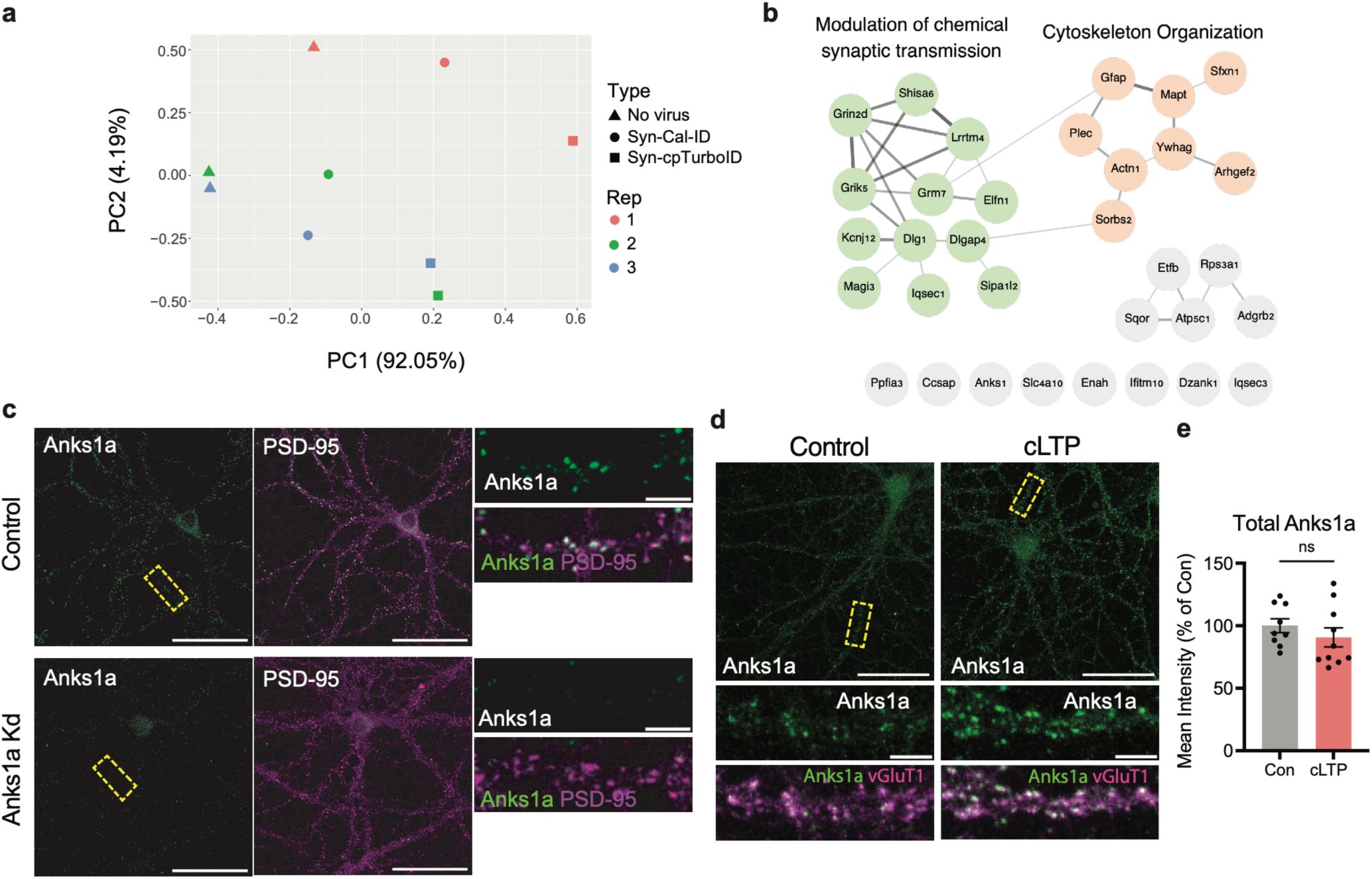
Proteomic screen using synaptic Cal-ID in vivo identifies Anks1a as a novel activity-dependent synaptic protein. a, PCA of mass spectrometry samples. b, STRING network analysis of top 7.6% of proteins ranked by enrichment ratio (confidence score cutoff = 0.25). Proteins cluster into Gene ontology (GO) biological processes including modulation of chemical synaptic transmission (FDR=0.0185) and cytoskeleton organization (FDR = 0.0069). Node colors indicate functional categories; grey nodes represent unclustered proteins. Edges represent protein–protein associations, with thickness and opacity scaling with STRING confidence score. c, Antibody validation for Anks1a antibody. d–e, Representative images (d) and quantification (e) of mouse primary neurons subject to cLTP. e, Mann-Whitney test, *n* = 9–10, 3 culture batches. Scale bars 50 μm (whole cell) or 5 μm (zoom). Data are presented as mean ± s.e.m. ns, not significant.

**Extended Data Figure 5.**
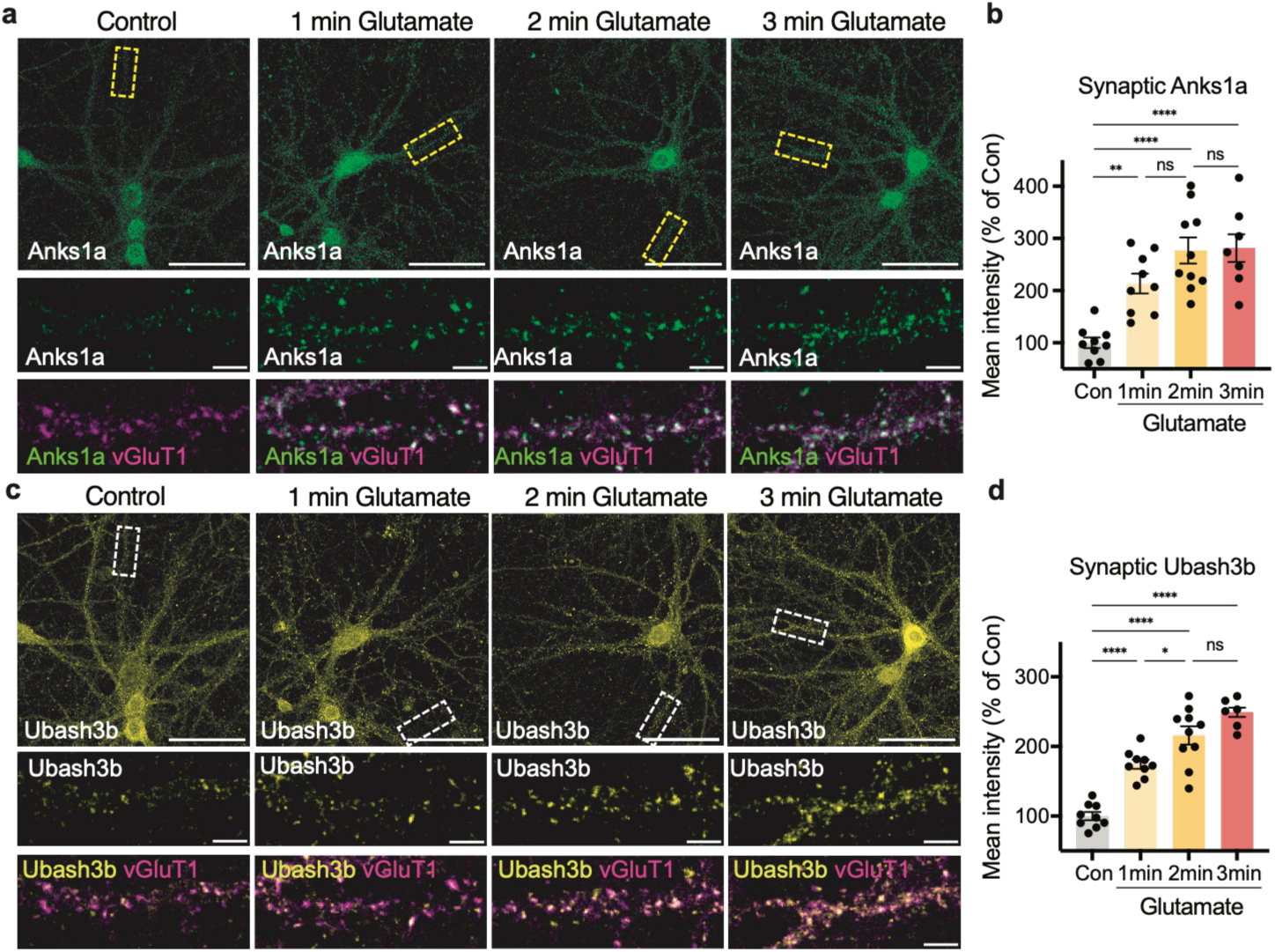
Time course of synaptic recruitment of Ubash3b and Anks1a. a–d, Representative images (a, c) and quantifications (b, d) of DIV19-21 mouse neurons following 15μM glutamate treatment over a time course. b, d, One way analysis of variance (ANOVA) with Sidak’s multiple-comparisons test, *n* = 8–10 cells, 3 culture batches. Scale bars 50 μm (whole cell) or 5 μm (zoom). Synaptic Anks1a/Ubash3b is defined as Anks1a/Ubash3b signal within vGluT1 or PSD-95 puncta. Data are presented as mean ± s.e.m. ns, not significant. *P < 0.05, **P < 0.01, ***P < 0.001, ****P < 0.0001.

**Extended Data Figure 6.**
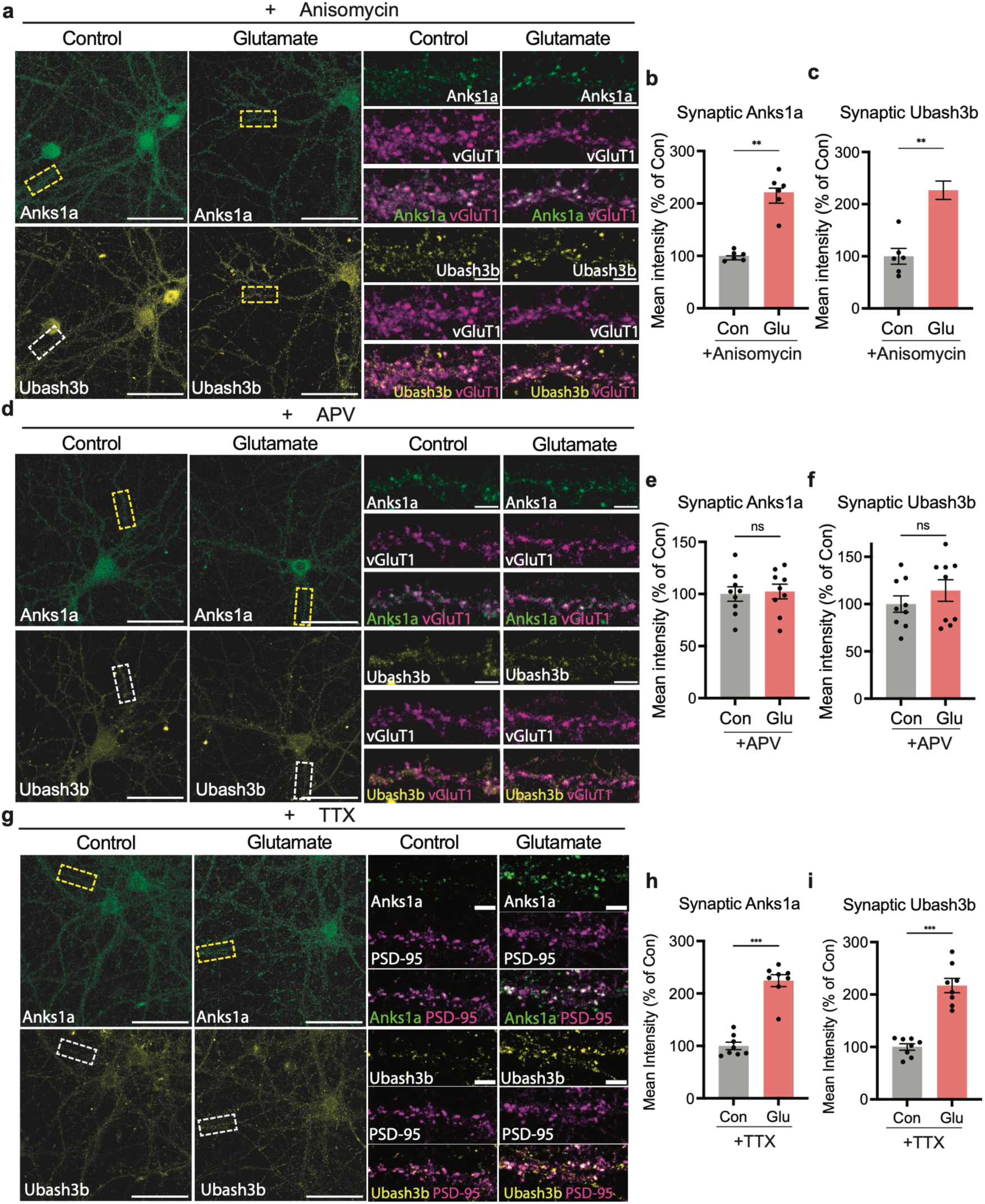
Ubash3b and Anks1a are rapidly recruited to synapses in a protein-synthesis independent and NMDAR-dependent manner. a–c, Representative images (a) and quantification (b, c) of mouse neurons treated with 15μM glutamate for 5 min in the presence of 25μM Anisomycin. b, c, Mann-Whitney test, *n* = 6 cells, 2 culture batches. d–f, Representative images (d) and quantifications (e, f) of immunostained mouse cultured neurons treated with 15μM glutamate for 5 mins in the presence of 50μM D-2-amino-5-phosphonovaleric acid (D-APV). Mann-Whitney test, *n* = 9 cells, 3 culture batches. Representative images (g) and quantifications (h, i) of immunostained mouse cultured neurons treated with 15μM glutamate for 5 mins in the presence of 1μM tetrodotoxin (TTX). Mann-Whitney test, n = 8, 2 culture batches. Synaptic Anks1a or Ubash3b is defined as Anks1a or Ubash3b signal within vGluT1 or PSD-95 puncta. DIV19–21 mouse primary cortical neurons. Scale bars 50μm (whole cell) or 5 μm (zoom). Data are presented as mean ± s.e.m. ns, not significant. **P < 0.01, ***P < 0.001,

**Extended Data Figure 7.**
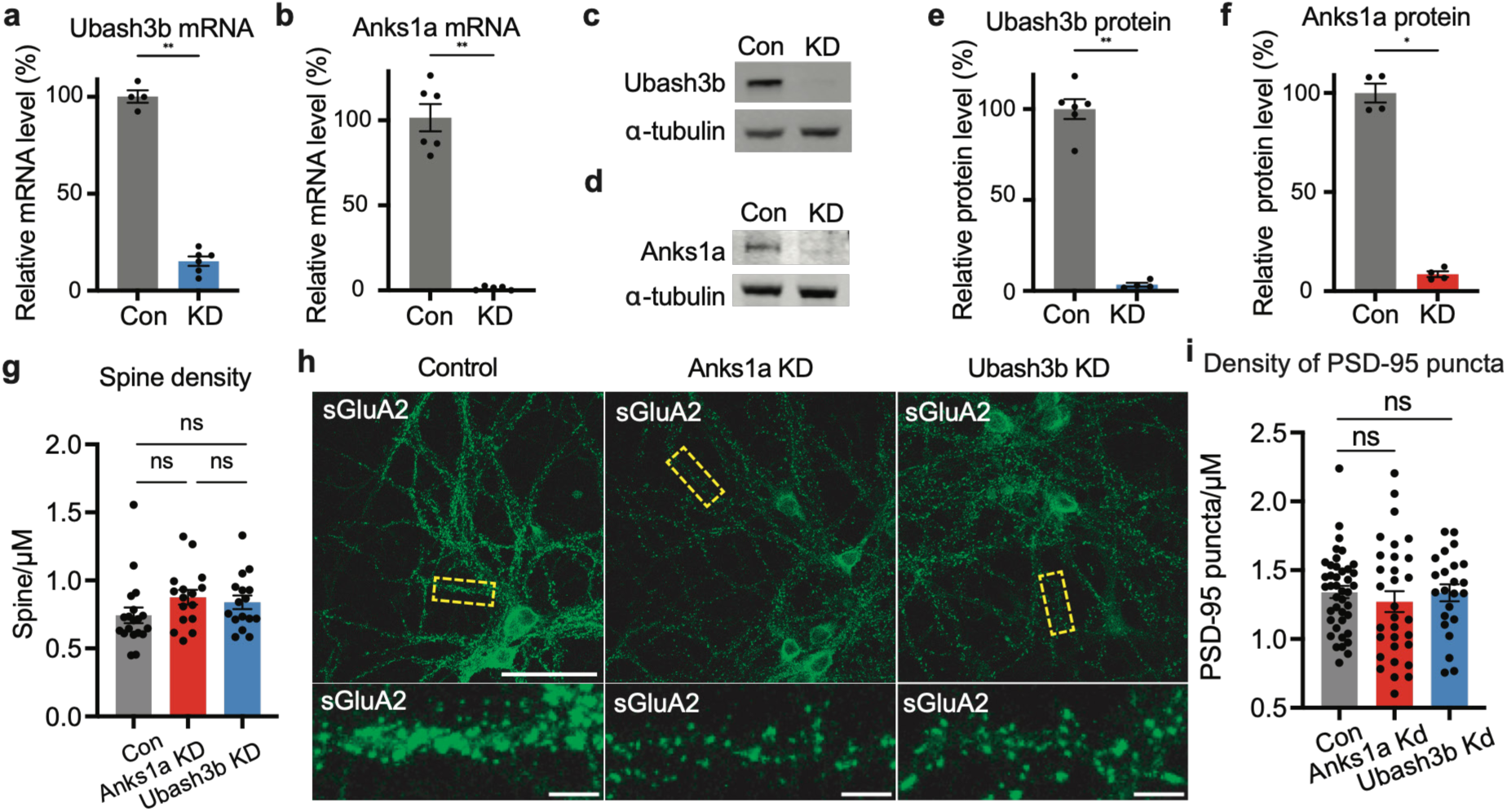
Loss of Anks1a or Ubash3b impairs synaptic structure and function. a, b, Quantification of qPCR results from Ubash3b-KD (a) or Anks1a-KD (b) DIV21 mouse primary neurons. Primary mouse neurons co-transduced with separate lentiviruses expressing dCas9 and gRNA. a, Mann-Whitney test, n = 4–6, Mann-Whitney test. b, Mann-Whitney test, n = 5–6. c–f, Representative western blot blots (c, d) and quantification from Ubash3b-KD (e) or Anks1a-KD (f) mouse primary neurons. e, Mann-Whitney test, n = 4–6. f, Mann-Whitney test, n = 4. g, Quantification of spine density from experiments related to Figure 5a. One-way ANOVA with Dunnett’s multiple-comparisons test, n =16-19 cells, 3 culture batches. h, Representative images from experiments related to Figure 5e. i, Quantification of PSD-95 puncta density from experiments related to Figure 5f. One-way ANOVA with Dunnett’s multiple-comparisons test, n = 23–41 cells from 2–3 culture batches. DIV19-21 primary mouse cortical neurons. Scale bars 50 μm (whole cell) or 5 μm (zoom). Data are presented as mean ± s.e.m. ns, not significant. *P < 0.05, **P < 0.01.

**Extended Data Figure 8.**
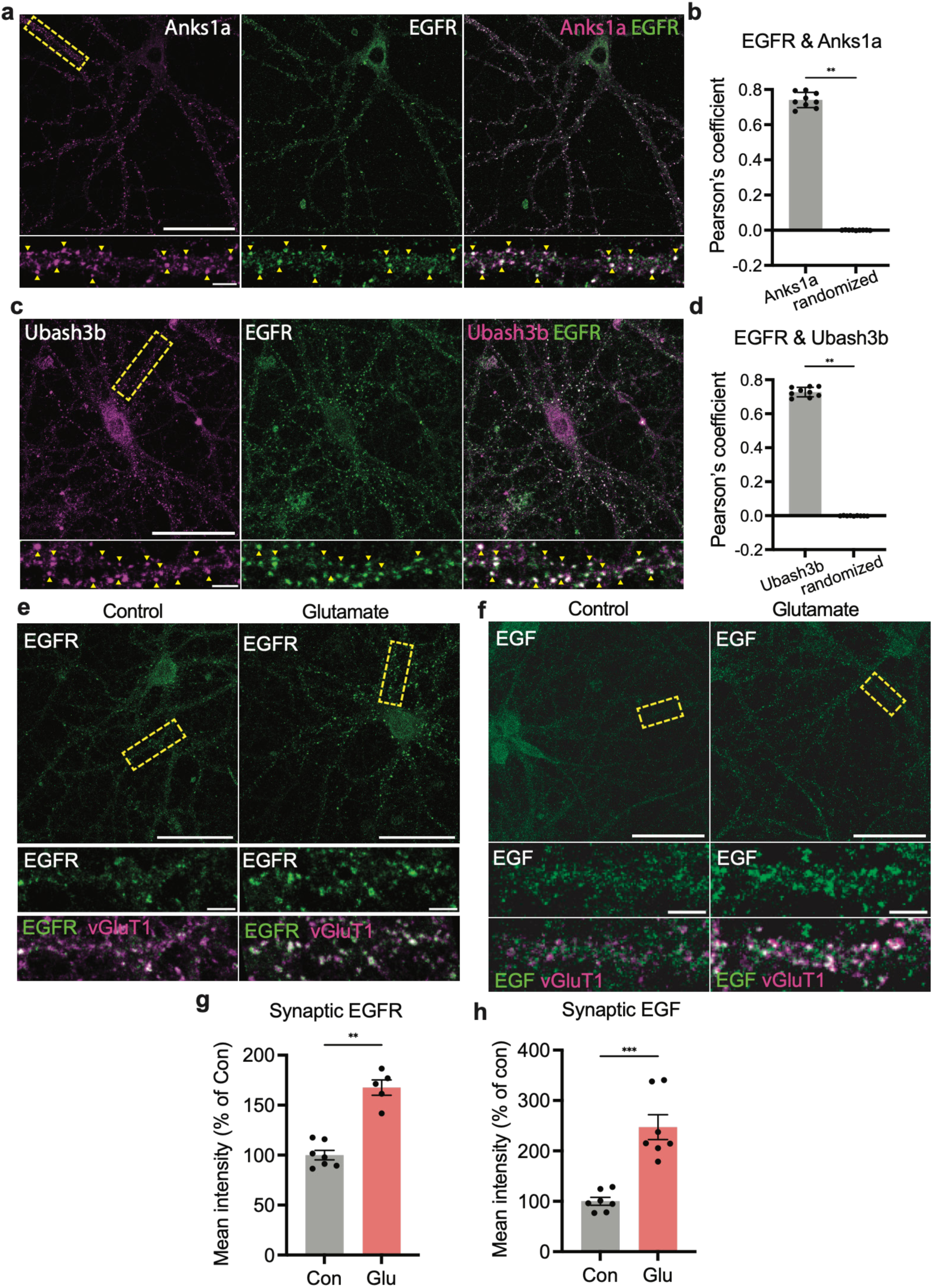
Activity-dependent EGFR accumulation at synapses requires Anks1a and Ubash3b. a–d, Representative images (a, c) and quantification (b, d) of immunostained primary mouse neurons. Yellow triangles indicate regions of colocalization. Wilcoxin matched-pairs signed rank test, n = 9 cells, 3 batches of cultures. e, h, Representative images (e) and quantification (g) of mouse neurons after glutamate treatment. Mann-Whitney test, n = 5–7, 2 culture batches. f, h, Representative images (g) and quantification (h) of mouse neurons after glutamate treatment. Mann-Whitney test, n = 7, 2 culture batches. DIV 19–21 primary mouse cortical neurons. Synaptic signal is defined as the signal within vGluT1 puncta. Scale bars 50 μm (whole cell) or 5 μm (zoom). Data are presented as mean ± s.e.m. **P < 0.01, ***P < 0.001.

**Extended Data Figure 9.**
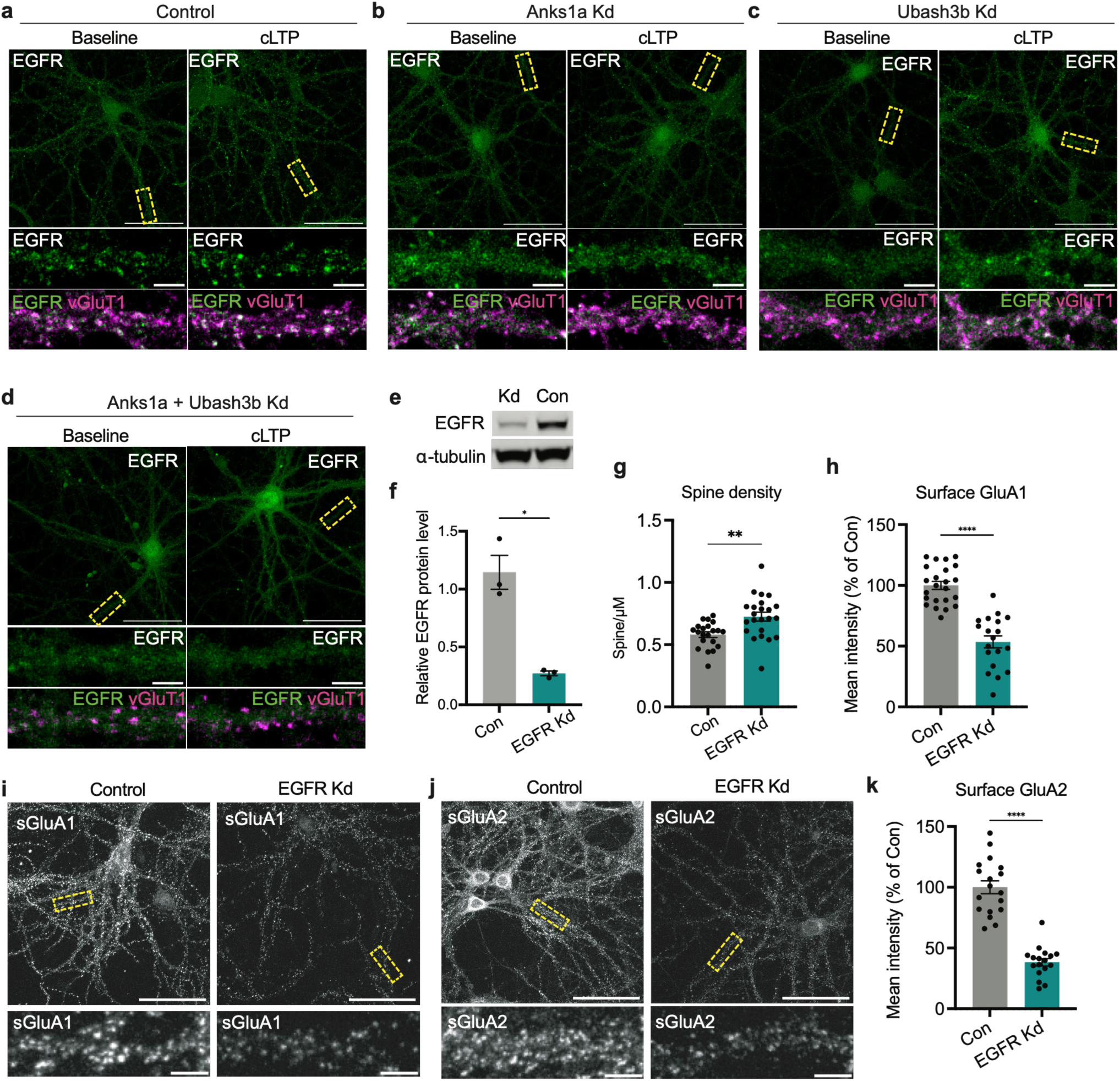
Loss of EGFR impairs synaptic structure and function. a–d, Representative images from experiments related to Figure 6e–h. e–f, Representative images (e) and quantification (f) of western blot from mouse neurons after CRISPRi knockdown of EGFR. Mann-Whitney test, n = 3. g, Quantification of spine density from Figure 6j. Mann-Whitney test, n = 21–23, 3 culture batches. h, i, Representative images (i) and quantification (h) of surface GluA1 in mouse neurons. h, Mann-Whitney test, n = 19-23, 4 culture batches. j, k, Representative images (j) and quantification (k) of surface GluA2 in mouse neurons. k, Mann-Whitney test, n = 17-18, 3 culture batches. DIV 21 primary mouse cortical neurons. Scale bars 50 μm (whole cell) or 5 μm (zoom). Data are presented as mean ± s.e.m. ns, not significant. *P < 0.05, **P < 0.01, ****P < 0.0001.

**Extended Data Figure 10.**
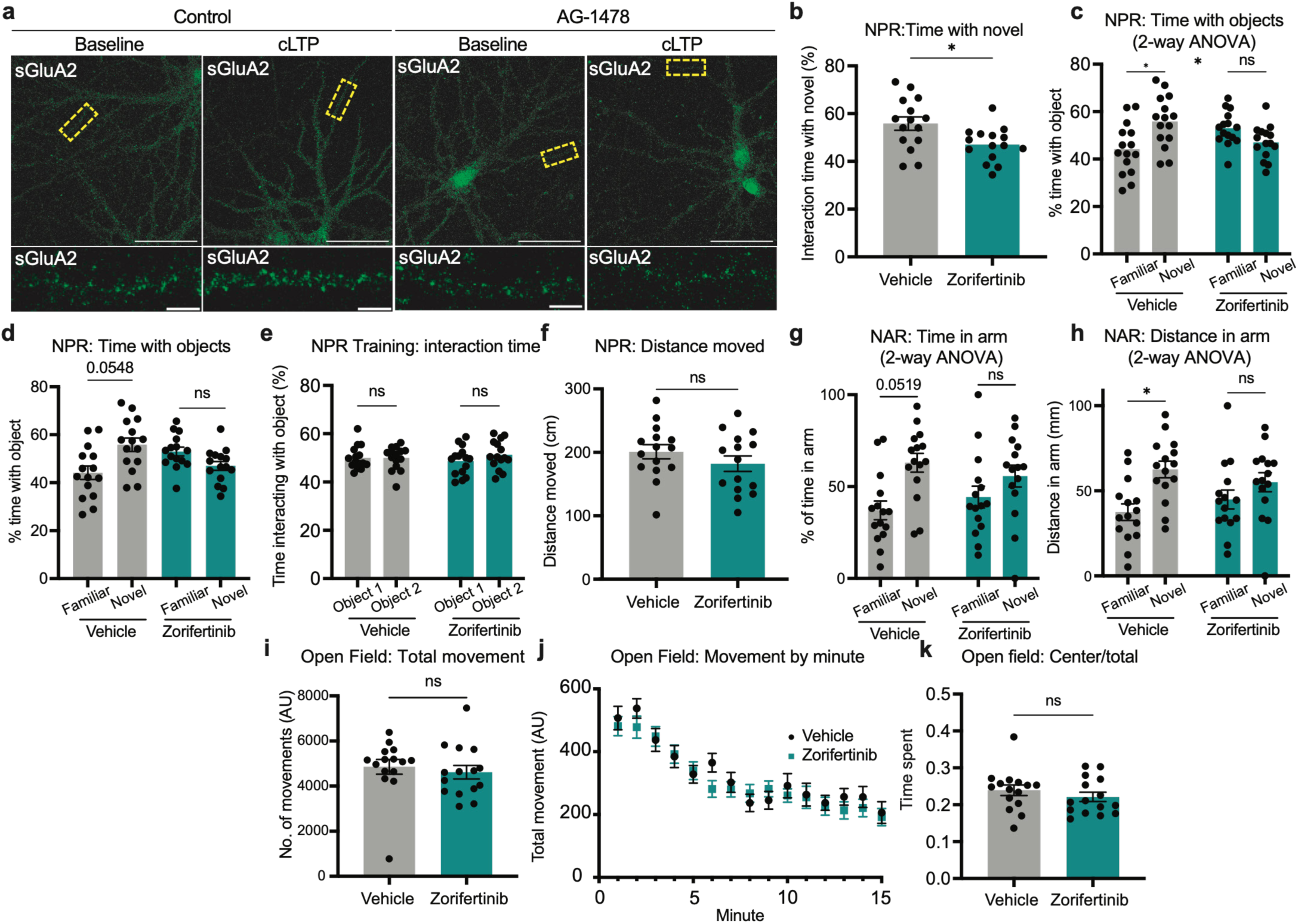
Acute EGFR inhibition impairs synaptic plasticity and learning and memory. a, Representative images of immunostained primary mouse neurons treated with AG-1478 or control and immunostained for surface GluA2 20 min after cLTP treatment. b–k, 4–5-month-old mice treated with vehicle or zorifertinib were subject to novel place recognition (b–f), novel arm recognition (g, h) and open field (i–m) tasks, *n* = 15 mice for each group. b, Percentage of time interacting with object in novel location, Welch’s t-test. c, Percentage time interacting with each object, two-way repeated-measures ANOVA, interaction significant. d, Percentage time interacting with each object, paired t-test. e, Percentage of time interacting with object in novel location, two-way repeated-measures ANOVA. f, Distance moved, Welch’s t-test. g, Percentage of time in each arm, two-way repeated-measures ANOVA, interaction ns. h, Distance in each arm, two-way repeated-measures ANOVA, interaction ns. i, Number of beam breaks, Mann-Whitney test. j, Beam breaks over time, two-way repeated measures ANOVA, interaction ns, all time points ns difference. k, Center beam breaks / total beam breaks, Welch’s t-test. Data are presented as mean ± s.e.m. ns, not significant. *P < 0.05.

